# Cancer systems immunology reveals myeloid—T cell interactions and B cell activation mediate response to checkpoint inhibition in metastatic breast cancer

**DOI:** 10.1101/2025.06.09.658361

**Authors:** Edgar Gonzalez, Jesse Kreger, Yingtong Liu, Xiaojun Wu, Arianna Barbetta, Aaron G. Baugh, Batul Al-Zubeidy, Julie Jang, Sarah M. Shin, Matthew Jacobo, Vered Stearns, Roisin M. Connolly, Won Ho, Juliet Emamaullee, Adam L. MacLean, Evanthia T. Roussos Torres

## Abstract

Sensitization of the immune-suppressed tumor microenvironment (TME) of breast cancer by histone deacetylase inhibition shows promise, but the mechanisms of sensitization are unknown. We investigated the TME of breast-to-lung metastases by combining experimental and clinical data with theory. Knowledge-guided subclustering of single-cell RNA-sequencing data and cell circuits analysis identified 39 cell states and salient interactions, of which myeloid, T cell and B cell subpopulations were most affected by treatment. Using functional immunologic assays, we verified that inhibition of the ICAM pathway partially recapitulated treatment effects. Mathematical modeling of tumor-immune dynamics confirmed that tumor reduction required simultaneous modulation of multiple TME interactions. We found evidence that treatment affected anti-tumor antibody production. Analysis of patient biopsies via spatial proteomics corroborated preclinical findings: in responders we observed increased B cell activation, mature tertiary lymphoid structures, and increased CD8+ T cell—macrophage distances with treatment. Overall, this study provides a framework for the discovery of cell-cell interactions that govern responses in complex TMEs.

**Statement of significance:** This study provides a framework for the discovery of cell-cell interactions that control responses in complex TMEs. We not only identify impactful tumor immunologic interactions that facilitate sensitization of the metastatic TME but also demonstrate how interdisciplinary data integration fuels cancer systems immunology to accelerate discovery of mechanisms of successful immunotherapeutic response in breast cancers and other previously unresponsive solid tumor types.

## INTRODUCTION

A major cause of death in patients with breast cancer is complications related to metastasis to distant organs^1^. Immune checkpoint inhibitors (ICIs) can stimulate anti-tumor T cell immunity by deactivating inhibitory signals in the tumor microenvironment (TME) and have been proven therapeutically effective. However, in the context of breast cancer, ICIs have been ineffective when used as single-agent therapy. Thus far, only one ICI is approved in combination with chemotherapy in specific cases of triple-negative breast cancer (TNBC), the most aggressive subtype for which there are no targeted therapies^2,3^. Numerous immune and stromal cells within the TME contribute to intrinsic resistance, including cancer-associated fibroblasts (CAFs), regulatory T cells (Tregs), macrophages, and myeloid-derived suppressor cells (MDSCs), all of which can be found in significant proportions in breast tumors and are associated with inhibition of anti-tumor immune responses and poor prognosis^4–7^.

In recent years, epigenetic agents have been studied with single-agent ICIs to modulate the immune response within the TME via their effects on host immune cells^8,9^. We and others have found that inhibition of class I histone deacetylases (HDACs) with the HDAC inhibitor (HDACi) entinostat (E), when given with anti-PD-1 (P) and anti-CTLA-4 (C), increases survival and inhibits breast-to-lung metastasis in mouse models of breast cancer^8,10^. These results supported our Phase Ib trial (NCI-9844), where we found that this same treatment combination resulted in a 25% objective response rate in patients with metastatic HER2-negative breast cancer (40% in patients with TNBC), demonstrating that EPC is also effective in patients^11^. Previous work has shown that changes to the immune TME associated with an anti-tumor response with EPC treatment, initiated by E, are intricate and complex, including M2-like to M1-like macrophage polarization, decreased suppression of CD8+ T cells by granulocytic-MDSCs (G-MDSCs), increased infiltration of CD8+ T cells, and lowered infiltration of G-MDSCs and Tregs, implicating suppressive immune cells and CD8+ T cells^8,10,12^. Entinostat can also affect cancer cells directly by stimulating apoptosis, cell cycle arrest, and immune editing of tumor antigens^13^. The genome-wide impact of entinostat on gene expression likely alters the complex crosstalk between cancer cells and immune cells, contributing to the treatment response observed in patients and preclinical models.

To unravel the complexities of global gene expression changes and subsequent effects on cellular crosstalk and immune function induced by entinostat within the TME, cancer systems immunology led us to integrate findings from our preclinical studies with mathematical modeling to uncover a response mechanism. We then investigated whether proposed mechanisms could be validated on patient samples from our treatment-matched clinical trial NCI-9844. This integrative approach yielded a powerful and highly efficient workflow culminating in mathematical models that inform mechanistic investigation.

## RESULTS

### Highly immune-infiltrated breast-to-lung metastases reveal tumor-immune responses to checkpoint inhibition

Given that treatment with HDACi with or without dual ICI is systemic and that many cell-cell interactions can contribute to response mechanisms, we sought a full characterization of the cellular components of breast-to-lung metastases. We utilized our recently described metastatic derivative NT2.5-LM^14^ cell line that forms spontaneous breast-to-lung metastases following injection into the mammary fat pad of NeuN mice. Mice were treated with entinostat (E) in combination with checkpoint inhibitors anti-PD-1 (P) and/or anti-CTLA-4 (C). We observed a slight yet significant improvement in survival in EPC-treated mice compared to mice treated with vehicle (V), which succumbed to symptoms of lung metastases by day 48 post-injection (**Figure 1A**). This contrasts with the 4T1 model of triple negative breast cancer, for which Kim *et al*. observed complete elimination of spontaneous lung metastasis in mice treated with EPC at 6 weeks post-treatment^8^. While this discrepancy could be attributed to differences in TME between the flank injection model used by Kim *et al*. and the mammary fat pad injection model of breast cancer metastasis used for NT2.5-LM, these results nevertheless support the use of the 4T1 model to mimic responders and the NT2.5LM model to mimic non-responders, as reported in our metastatic breast expansion cohort of NCI-9844^9^. To characterize cellular components of metastases, we performed single-cell RNA sequencing (scRNA-seq) of parental NT2.5 breast tumors^12^ and NT2.5-LM breast-to-lung metastases treated with different combinations of entinostat, anti-PD-1, and anti-CTLA-4 for a total six treatment groups. We compared 54,636 cells in mammary primary tumors with 58,097 sequenced cells isolated from macrodissected lung metastases. Following quality control, filtering, and batch correction, we identified cell states via Louvain clustering for the primary tumor and the lung metastases, producing six and eight clusters, respectively (**Figure 1B-C**). Both datasets contained clusters representing cancer cells, T cells, mature myeloid cells, and immature myeloid-derived suppressor cells (MDSCs). Expression of genes representative of each cell state supported the classification of these cells (**Figure 1D-E**, **Supplementary File 1**). Both datasets also contained endothelial and fibroblast populations, although fibroblasts differed between organs. In the breast microenvironment, fibroblasts expressed genes associated with cancer-associated fibroblasts (CAFs)^15^, whereas in the lung, the cells closely resembled lipofibroblasts^16^ by transcriptional state (**Figure 1D, 1E, Figure S1**). We also identified B cells and Natural Killer (NK) cells in lung metastases, unique immune populations that were not detectable in the primary breast tumor microenvironment. Investigation into the effect of treatment on the cellular composition of the two tumor types at this level revealed no statistically significant shifts in cell state composition in either breast tumors (**Figure 1F**) or lung metastases (**Figure 1G**).

**Figure 1.**
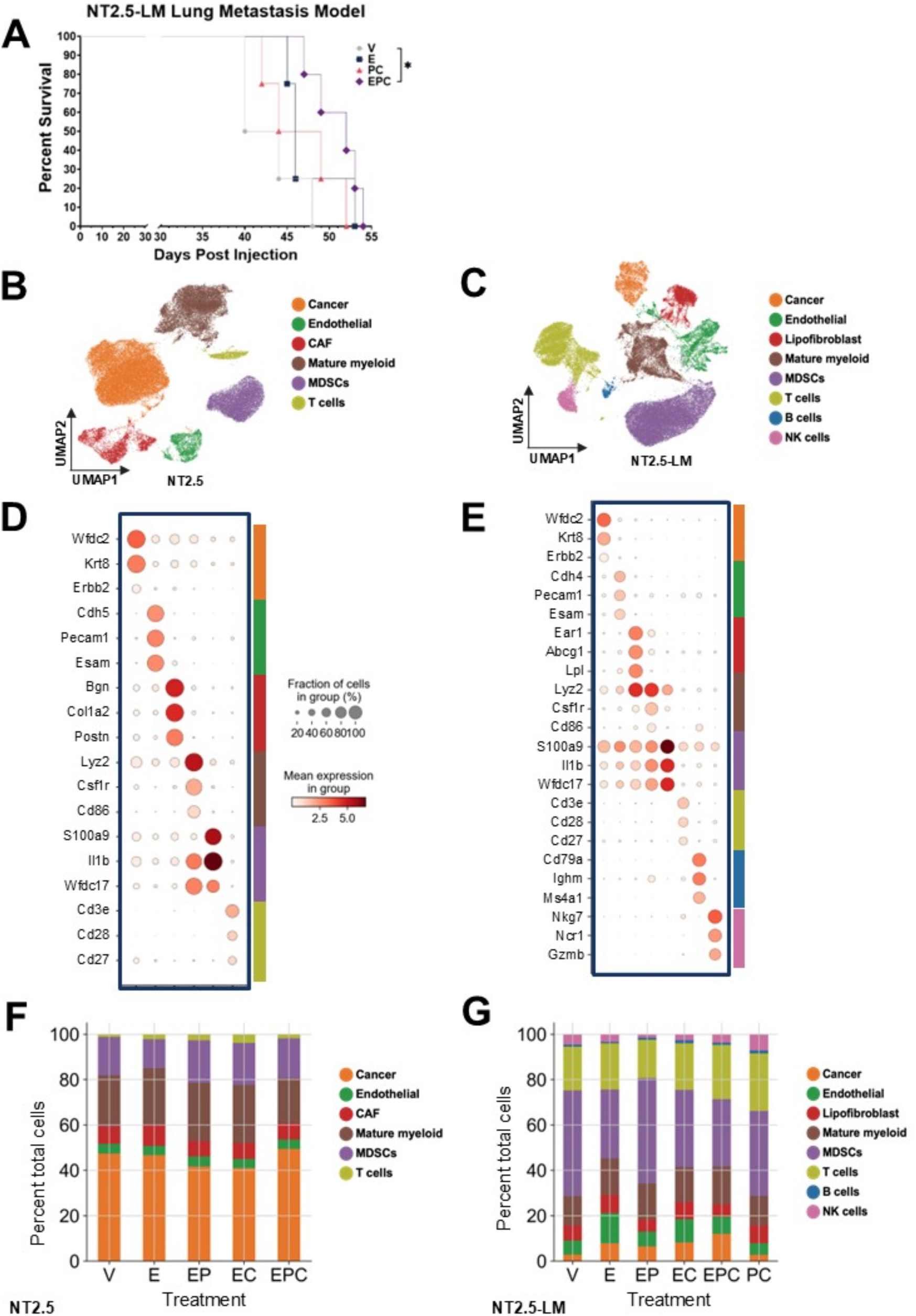
Survival and major cell populations in the NT2.5-LM breast-to-lung metastasis model and NT2.5 primary tumor model with entinostat and checkpoint inhibitors. **A.** NeuN mice (n=4-5/group) were injected in the tail vein with 100,000 NT2.5-LM cells and treated with a combination of 5 mg/kg entinostat (E) 5 times/week, 100 µg/dose anti-PD-1 (P) twice/week, and 100 µg/dose anti-CTLA-4 (C) twice/week starting on day 24 post-injection of NT2.5-LM cells. Mice were euthanized as they developed symptoms of lung metastasis. Statistically significant differences were determined using the logrank test for survival; *p<0.05. **B.** UMAP showing all major cell states identified in NT2.5 mammary tumors (n=4 mice/group); data from^12^. **C.** UMAP showing all major cell states identified in NT2.5-LM lung metastases (n=3-5 mice/group) via single-cell RNA-sequencing. **D-E.** Dot plots showing expression of signature marker genes in each major cell type identified in NT2.5 mammary tumors **(D)** and NT2.5-LM lung metastases **(E)** as detailed in supplementary table 1. **F-G.** Bar plots showing proportions of major cell populations identified in NT2.5 mammary tumors **(F)** and NT2.5-LM lung metastases **(G)** via scRNA-seq with treatments.

### Knowledge-guided subclustering of immune cell subsets identifies functionally relevant immune cell infiltration

To investigate heterogeneity within immune cell populations, we subjected each high-level immune cluster within NT2.5-LM lung metastases (**Figure 1C**) to unsupervised clustering and differential gene expression analysis. Subclusters that failed QC (high doublet scores, low gene counts, or high mitochondrial gene expression) were discarded, and immune cell states were assigned based on the gene expression in each subcluster, informed by a review of the literature (**Supplementary Table 1, Supplementary File 1**). Within the mature myeloid population, we identified 12 distinct cell states, including four macrophage, classical/non-classical monocyte, three neutrophil, and three dendritic cell states (**Figure 2A-B**). We identified a *C1qa/b/c*+ macrophage population with gene expression matching the C1q macrophage subtype observed in various preclinical models and human tumors^17^, along with a closely related metabolically activated macrophage^18–20^ population with high expression of genes involved in lipid metabolism (*Cd36*, *Lpl,* and *Plin2*), lysosomal function (*Ctss*, *Lamp2*, and *Lipa*), and inflammation (*Il18)* (**Figure S2A**). Both macrophage subtypes have been described as pro-tumoral, and C1q macrophages are associated with a poor prognosis. We also identified proliferating macrophages (*Cdk1+*) and a *Ccr2*+ macrophage population expressing high levels of classical monocyte markers, as described in other tumor models^21,22^. Three neutrophil subclusters and three DC cell states were identified (**Figure S2B-C**), with the TAN1 and TAN2 neutrophil cell states expressing notably high levels of pro-tumor genes: *Fn1* and *Ccl9* in TAN1 and *Lcn2*, *Mmp8*, and *Mmp9* in TAN2^15,23–25^. Treatment with entinostat (E) resulted in increased proportions of C1q macrophages (false detection rate [FDR] of 0.008), metabolically activated macrophages (FDR of 0.003), and conventional dendritic cells (cDC1 with FDR of 0.07 and cDC2 with FDR of 0.007) (**Figure 2E and S2D**). None of the shifts observed with E were preserved with EPC, although there was a possible increase in C1q macrophages that did not meet significance (FDR of 0.14).

**Figure 2.**
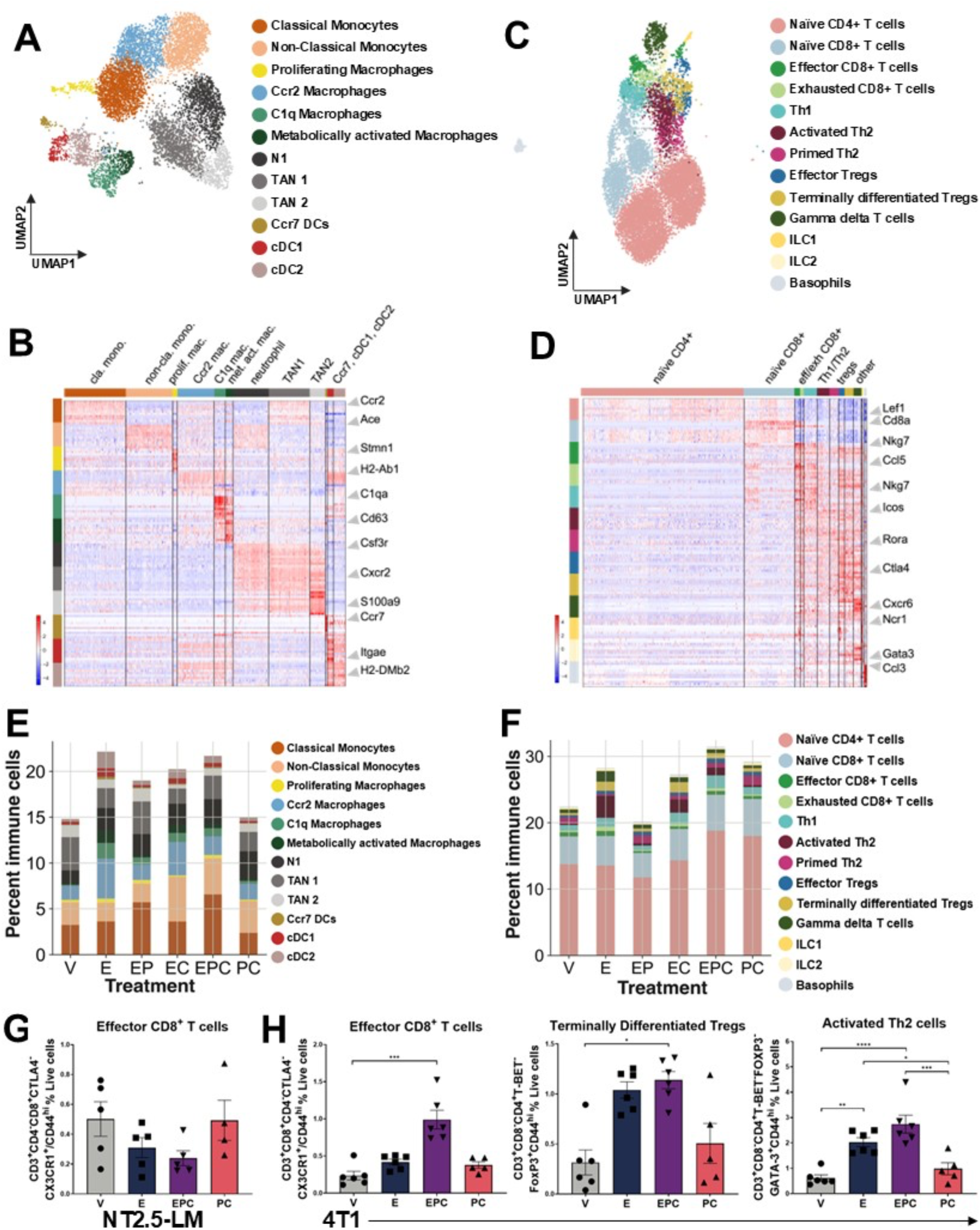
Treatment effects on mature myeloid and T cell proportions in NT2.5-LM via scRNA-seq and CD8+ T cell proportions in NT2.5-LM and 4T1 via flow cytometry. **A.** UMAP of subclusters identified in the mature myeloid cell major cluster in NT2.5-LM lung metastases via scRNA-seq (Figure 1C). **B.** Heatmap with z-scores demonstrating the expression of top differentially expressed genes observed in the mature myeloid cell populations identified in **(A)**. Arrows point out signature marker genes for each cell state in panels B and D. **C.** UMAP of subclusters identified in the T cell major cluster (Figure 1C) in NT2.5-LM lung metastases via scRNA-seq. **D.** Heatmap with z-scores demonstrating the expression of top differentially expressed genes observed in T cell populations identified in C. **E-F.** Stacked barplots showing the proportions of mature myeloid cell populations (E) or T cell populations (F) identified in NT2.5-LM lung metastases with treatments. **G-H.** Flow cytometry analysis of cell proportions, shown as a percentage of total single, live cells for T cell subsets matching T cell populations identified via scRNA-seq in NT2.5-LM (G) or 4T1 (H) lung metastases with treatments. Antibody panels used for flow cytometry are found in Supplementary File 2. *p<0.05, **p<0.01, ***p<0.001, ****p<0.0001. V = vehicle, E = entinostat, P = anti-PD-1, C = anti-CTLA-4. n=4-5/group for G and n=5-6/group for H. Statistically significant differences were determined using the Kruskal-Wallis test for effector CD8+ T cells and the one-way ANOVA for activated Th2 cells and terminally differentiated Tregs in panels G-H.

T cells within the lung metastases were subclustered into 13 distinct cell states. Of these, naïve CD4+ T cells and naïve CD8+ T cells were the most abundant, with smaller populations of effector CD8+ T cells, regulatory T cells (Tregs), T helper cells (Th1/Th2), and gamma-delta T cells (ɣδ T cells) (**Figure 2C-D**). In total, three cell states were identified for CD8+ T cells (naïve, effector, and exhausted CD8+ T cells) and six for CD4+ T cells (Th1, primed Th2, exhausted Th2, naïve CD4+ T cells, effector Tregs, and terminally differentiated Tregs) based on characteristic expression of signature marker genes (**Figure S3A-B**). Although primed Th2 cells have not been previously described in the literature, we identified a cell state with high levels of naïve T cell marker *Ccr7* and early activation marker *Cd69*, distinguishing them from activated Th2 cells with higher expression of the *CD44* activation marker^26–28^. The remaining subclusters were designated as ɣδ T cells, ILC1, ILC2, and basophils based on characteristic gene expression (**Figure S3C**). We used ProjectTILs^29^ to validate the cell type labels assigned to T cell states and observed high concordance between ProjectTIL assignments and our supervised T cell state assignments (**Figure S3D**). Treatment with entinostat resulted in significant increases in the proportions of activated Th2 cells (FDR of 0.008) and terminally differentiated Tregs (FDR of 0.038) (**Figure 2F, S3E table**). Although no significant shifts in cell proportions were observed with EPC, the shift from primed Th2 cells with V to activated Th2 cells with E was maintained with EPC.

To assess concordance between the frequencies of immune cell subtypes observed by scRNA-seq (**Figure 2E-F**) vs. antibody-based sorting, we analyzed cells from NT2.5LM breast-to-lung metastases following treatment with V, E, PC, and EPC via multi-parametric flow cytometry. Similar proportions of mature myeloid and T cell populations were observed (**Figure 2G, S2E, S3F**). The increases in subtypes of mature myeloid cells and T cells predicted to occur with E by scRNA-seq were not recapitulated by flow cytometry. The same gating strategies were used to analyze immune cell subtypes within lung metastases obtained from mice using the 4T1 model. As observed with NT2.5-LM, there were no significant shifts in mature myeloid cell subtype proportions with treatment (**Figure S4**). In agreement with the NT2.5-LM scRNA-seq data, there was a significant increase in the proportion of activated Th2 cells with E, which was maintained with EPC (p<0.0001). (**Figure 2H, S5**). The proportion of effector CD8+ T cells also increased with EPC, highlighting a potentially synergistic effect in the 4T1 model between E and PC (**Figure 2H**).

We identified two cell states within the MDSCs: monocytic (M)-MDSCs and granulocytic (G)- MDSCs (**Figure S6A-B**). Treatment with E significantly decreased the number of G-MDSCs (FDR<0.05) (**Figure S6C table**). As previously observed for NT2.5 primary tumors, analysis of NT2.5-LM lung metastases by flow cytometry revealed no significant changes in M-MDSC or G-MDSC proportions with treatment (**Figure S6D**). Similarly, no significant changes were observed in the infiltration of M-MDSCs or G-MDSCs into lung metastases in the 4T1 model with treatment (**Figure S6E**). We found evidence for three subpopulations within the NK cell major cluster, which were identified as mature NK, immature NK, and tumor-primed NK cells based on expression of signature marker genes such as *Gzma*, *Cd27*, and *Ifng*, respectively (**Figure S6F-G**). No significant differences in proportions of NK cell states were observed with treatment, although very few NK cells were observed overall with EP (**Figure S6H**). Analysis of NK cell infiltration via flow cytometry revealed no differences in NK cell proportions with treatments compared to V (**Figure S6I**).

### Combination therapy increases B cell infiltration and promotes the generation of cancer cell-targeting antibodies

The B cell cluster, the smallest among the major cell clusters, consisted of 4 B cell populations, plasma cells, and plasmacytoid dendritic cells (pDCs), with notable expression of signature marker genes (**Figure 3A-B**). The four B cell populations included naïve B cells, Bregs, and two mature B cell populations, distinguished by higher expression of memory B cell markers *Cd37* and *Cr2*^30^ in the mature B cells 2 subtype. Treatment with EPC increased the proportion of mature B cell 2 and plasma cells, and both E and EPC increased the ratio of mature B cells 2 to mature B cells 1 (**Figure 3C**). Plasma cell proportions also increased with EPC compared to E, but not with PC, suggesting a synergistic effect between E and PC. There were no treatment effects on total B cell infiltration by flow cytometry, concordant with scRNAseq results (**Figure 3D**). However, CD11b^+^ B cell proportions increased in mice treated with EPC (∼15% versus ∼3% for vehicle), matching the increase in CD11b^+^ plasma cells observed with EPC treatment in the scRNA-seq dataset (**Figure 3E, S7A**). Expression of various MHC-II genes, including *H2-Aa* and *H2-Ab1,* increased in the mature B cell 2 and Breg populations with treatments (**Figure 3F**). Cell-cell communication network inference with CellChat^31^ revealed increased signaling to primed Th2 and exhausted CD8+ T cells from mature B cells 2, Bregs, and plasma cells with EPC (**Figure S7B).** EPC decreased predicted cell-cell interactions between Bregs/mature B cells 2 mediated via ligand-receptor (LR) pair *Cd22---Ptprc*, the expression of which associates with the inhibition of B/T cell receptor signaling^32,33^ (**Figure 3G**). EPC treatment also decreased the predicted interaction between Bregs and exhausted CD8+ T cells via *Cd274---Pdcd1,* and increased the predicted interaction between mature B cells 2 and primed Th2 cells via *Cd86-Cd28*, which is crucial for optimal activation of Th2 cells and IL-4 production^34^ (**Figure S7C-D**).

**Figure 3.**
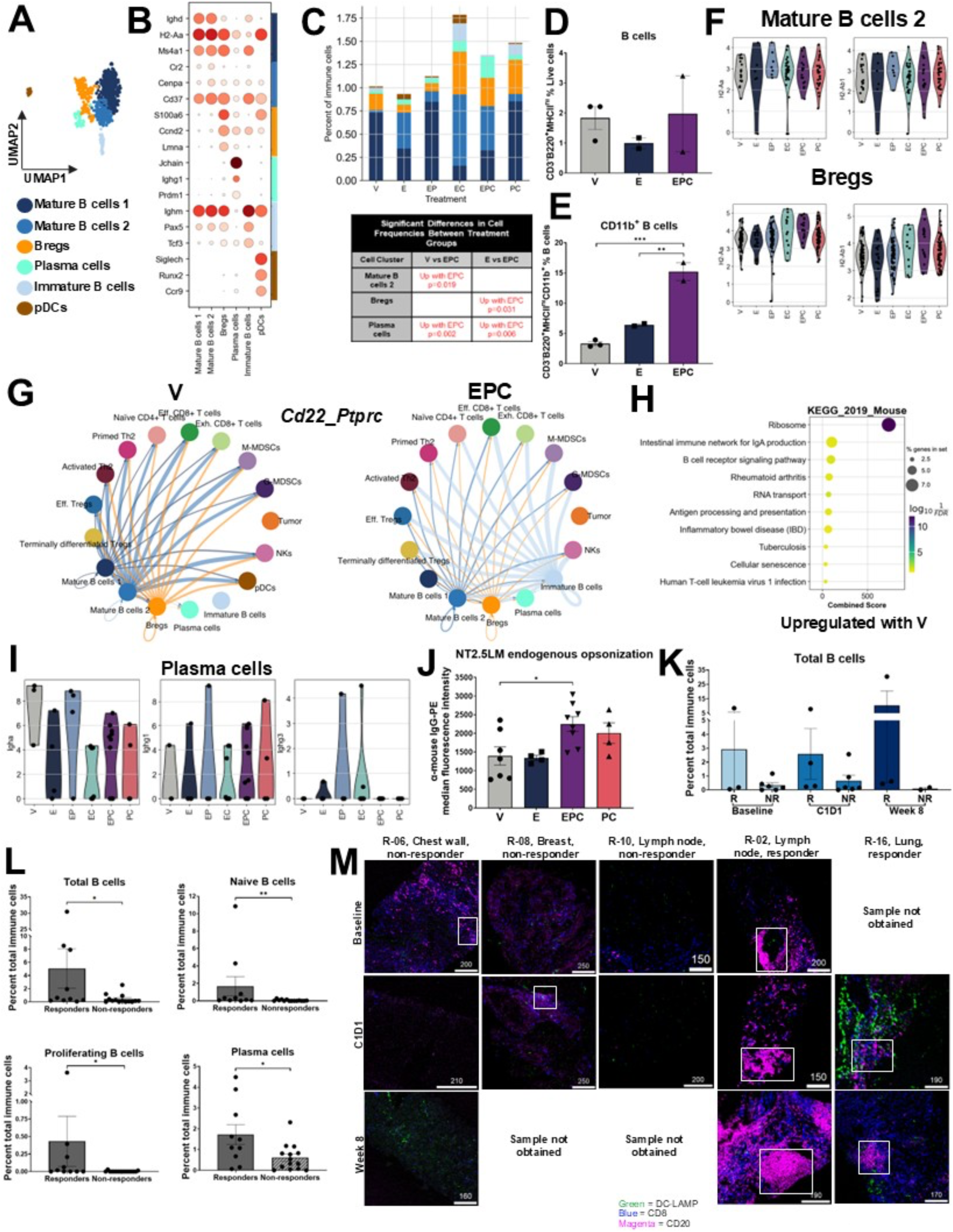
Analysis of treatment effects on B cells in NT2.5-LM lung metastases using scRNA-seq and flow cytometry and B cells in patient tumors using spatial proteomics. **A.** UMAP of subclusters identified in the B cell major cluster (Figure 1C) in NT2.5-LM lung metastases via scRNA-seq. **B.** Dot plot showing the expression of signature markers for the B cell, plasma cell, and pDC populations identified in A. **C.** Stacked bar plots showing B cell, plasma cell, and pDC proportions with treatments and accompanying table with statistically significant differences identified between treatment groups. **D-E.** Flow cytometry analyses of NT2.5-LM lung metastases showing proportions of total B cells as a percentage of total single, live cells and CD11b+ B cells as a percentage of total B cells with V, E, and EPC treatments (n=4-5/group). V = vehicle, E = entinostat, P = anti-PD-1, C = anti-CTLA-4. **p<0.01, ***p<0.001. **F.** Violin plots showing the expression of *H2-Aa* and *H2-Ab1* in mature B cells 2 and Bregs subclusters with treatments. **G.** Circos plots depicting the *Cd22*_*Ptprc* interactions between cell populations included in the cell chat analysis for B cells. Arrows point from cellular sources of *Cd22* to cells interacting with *Cd22* via *Ptprc* (CD45), with increased arrow thickness representing increased interaction strength. **H.** KEGG analysis (2019 version) of pathways upregulated in vehicle-treated mice compared to EPC in total B cells/plasma cells from NT2.5- LM lung metastases. **I.** Violin plots showing the expression of *Igha, Ighg1,* and *Ighg3* in plasma cells with treatments. **J.** Flow cytometry analysis of HER2+ cancer cells (gated as CD45-HER2+ single, live cells) from NT2.5-LM lung metastases obtained from treated mice (n=4-8/group) using a PE-conjugated anti-mouse IgG to detect levels of endogenously bound IgG via quantification of median fluorescence intensity. **K.** Spatial proteomics analysis of patient tumor tissues stratified by treatment response and timepoint showing the proportion of total B cells (n=2-6/group). R = responder. NR = Non-responder, C1D1 = post-entinostat, week 8 = post entinostat + anti-PD-1 + anti-CTLA-4. J. **L.** Proportions of total B cells, proliferating B cells, naïve B cells, and plasma cells shown in responders compared to non-responders, including samples from all timepoints (n=10-14/group). *p<0.05, **p<0.01. **M.** Visualization of raw signals for CD8 (blue, CD8+ T cells), CD20 (magenta, B cells), and DC-LAMP (green, mature DCs) in pre-processed IMC images of tumor tissues for patients R-06, R-08, R-10, R-02, and R-16 (left to right) at baseline, C1D1, and week 8 timepoints. White rectangles within images outline tertiary lymphoid structures (TLS) or germinal centers (in lymph nodes). Samples were not obtained for patients R-08 and R-10 at week 8 and R-16 at baseline. V = vehicle, E = entinostat, P = anti-PD-1, C = anti-CTLA-4. Statistically significant differences between cell proportions in panel C were determined via unpaired t-tests. Statistically significant differences were determined using the one-way ANOVA for panels E and J and the Mann-Whitney test for panel L. Antibody panels used for flow cytometry are found in Supplementary File 2.

KEGG pathways analysis on total B cells uncovered a reduction in the intestinal immune network for IgA production with EPC (p_adj_=0.03; **Figure 3H**). In addition, PC and EPC treatment decreased *Igha* expression, increased *Ighg1* expression, and had no effect on *Ighg3* or *Ighg2a* expression in plasma cells, which remained at low or undetectable levels (**Figure 3I**). Altogether, these results indicate that EPC induced a class switch recombination from predominant production of IgA to IgG1. To determine functional implications, NT2.5-LM cells were incubated with serum from EPC-treated mice, resulting in significantly increased opsonization compared to V (**Figure S7E**). Flow cytometry of HER2+ cancer cells isolated from lung metastases also demonstrated increased endogenous opsonization with EPC treatment (**Figure 3J**).

To validate preclinical findings and examine changes in immune cell infiltration and interactions in patients, we performed spatial proteomics using imaging mass cytometry (IMC) on tumor biopsies from our trial, NCI-9844^9^, in which patients with metastatic breast cancer were treated with entinostat (E), nivolumab (anti-PD-1, P), and ipilimumab (anti-CTLA-4, C). Hormone receptor positive (HR+, n=5) and TNBC (n=6) tissues from breast tumors and metastases obtained from chest wall, liver, lung, and lymph node (LN) were collected at the time of trial enrollment (baseline), after 2 weeks of E (C1D1), and after eight weeks of EPC (week 8). Five patients were considered clinical responders according to RECIST criteria, and 6 patients were considered non-responders^3^. We found increased total B cell and plasma cell proportions in responders across all timepoints (**Figure 3K, S7F**). When combining all timepoints, proportions of total B cells, plasma cells, proliferating B cells, and naive B cells were all higher in responders (**Figure 3L**). Since B cells and plasma cells are associated with the presence of tertiary lymphoid structures (TLS) in tumors^35^, we assessed samples for clustering of B cells/plasma cells, indicating the presence of a TLS. Samples that had a detectable TLS or germinal center (lymph nodes) showed increased proportions of plasma cells, activated B cells, and proliferating B cells (**Figure S7G**). Early-stage TLS^36^ were found in three patient samples at baseline and C1D1, along with a mature TLS with follicular morphology in a responding patient at week 8 (**Figure 3M, S7H**). In addition, there were observable follicular structures in all lymph node biopsies from a responder but none in a non-responder. Functional marker analysis revealed increased expression of the survival/proliferation marker CD137^37,38^ in total B cells and decreased expression of the LAG3 immunosuppression marker^39^ in mature B cells in responders treated with EPC (**Figure S7I**). HLA-DR expression was also significantly higher in mature B cells at baseline and week 8 in responders, which is associated with increased antigen presentation^40^.

### Combination therapy shifts cancer cells toward a more mesenchymal and stem-like state

We investigated cancer cell heterogeneity within this syngeneic mouse model, which we predicted would be low, given the clonal nature of the NT2.5-LM cell line. Unsupervised subclustering of cancer cells identified three cell states, none of which were altered by treatment (**Figure S8A-B**). Differential gene expression between cancer cell states revealed little variation overall (**Figure S8C)**: cell cycle genes were upregulated in tumor state 2, and ribosomal genes and S100a8/9 were upregulated in tumor state 1. Analysis of cell cycle activity revealed tumor state 2 to contain mostly actively cycling cells (G2M or S phase), while tumor state 0 contained non-cycling cells (in G1 phase) primarily (**Figure S8D**). No significant changes were observed in cell cycle activity with treatment (**Figure S8E**).

Given the lack of evidence for cancer cell state heterogeneity driven by factors other than cell cycle or ribosomal activity, we hereafter considered the cancer clusters as a single cell state. To analyze treatment effects, we performed linear discriminant analysis to assess the extent to which treated cancer cells could be distinguished from untreated cells globally by their transcriptional state. Projecting cells along the first linear discriminant (LD1), E-treated cancer cells were most distinguishable transcriptionally from V (Earth mover’s distance (EMD) = 1.64), whereas PC-treated cells changed the least (EMD = 0.29), and EPC-treated cells were intermediate (EMD = 1.22) (**Figure S8F**). Comparison of differentially expressed genes with genes contributing to LD1 for each treatment comparison identified overlapping genes, such as S100a8/9, whose expression was consistently higher in V compared to any treatment. It also identified candidate genes specific to treatment, such as *Col11a1* with E (**Figure S8G**). We applied simple and iterative logistic regression to predict differences between treated and untreated cells. Top weighted genes included *Cd81* (PC) and *Vim* (EPC), the latter suggesting shifts in EMT-associated phenotypes with treatment (**Figure S9A**). Analysis of stemness and mesenchymal gene set scores, as well as PAM50 subtypes, uncovered a change towards a more stemness/mesenchymal phenotype with E, which was maintained with EPC for mesenchymal scores (**Figure S9B-C**). While no significant differences were observed in PAM50 basal scores, PAM50 myoepithelial scores significantly increased with treatment E and EPC (**Figure S9D**).

### Cell circuit analysis identifies the specific T cell---myeloid cell interactions that are most important for response

To identify prominent cell-cell interactions in the lung TME and study how they are affected by treatment, we inferred cell-cell communication networks from scRNA-seq using CellChat. Among the eight high-level cell types in the TME, cancer cells, lipofibroblasts, mature myeloid cells, and MDSCs were the largest sources of signaling (**Figure 4A**). However, analysis of the cell-cell communication at this level does not offer a complete picture: we need to investigate it at the level of the specific immune subtypes identified. For the 39 subclusters identified across eight major cell types, this would require analysis of 741 networks for pairwise networks (9,139 networks for triplets of cell states). Given this complexity, we developed an approach to reduce the dimensionality and investigated cell-cell signaling between important cell subsets via two comparisons. The first comparison (CC-suppression) was compiled from cell subtypes described as immunosuppressive in the literature and subtypes that are known to interact with these populations (**Figure 4B**). The second comparison, CC-model, was compiled to assess the treatment-relevance of our previously developed tumor-immune interaction model^41^, which discovered that interactions between NK cells, CD8+ T cells, and MDSCs are crucial in determining tumor growth outcomes.

**Figure 4.**
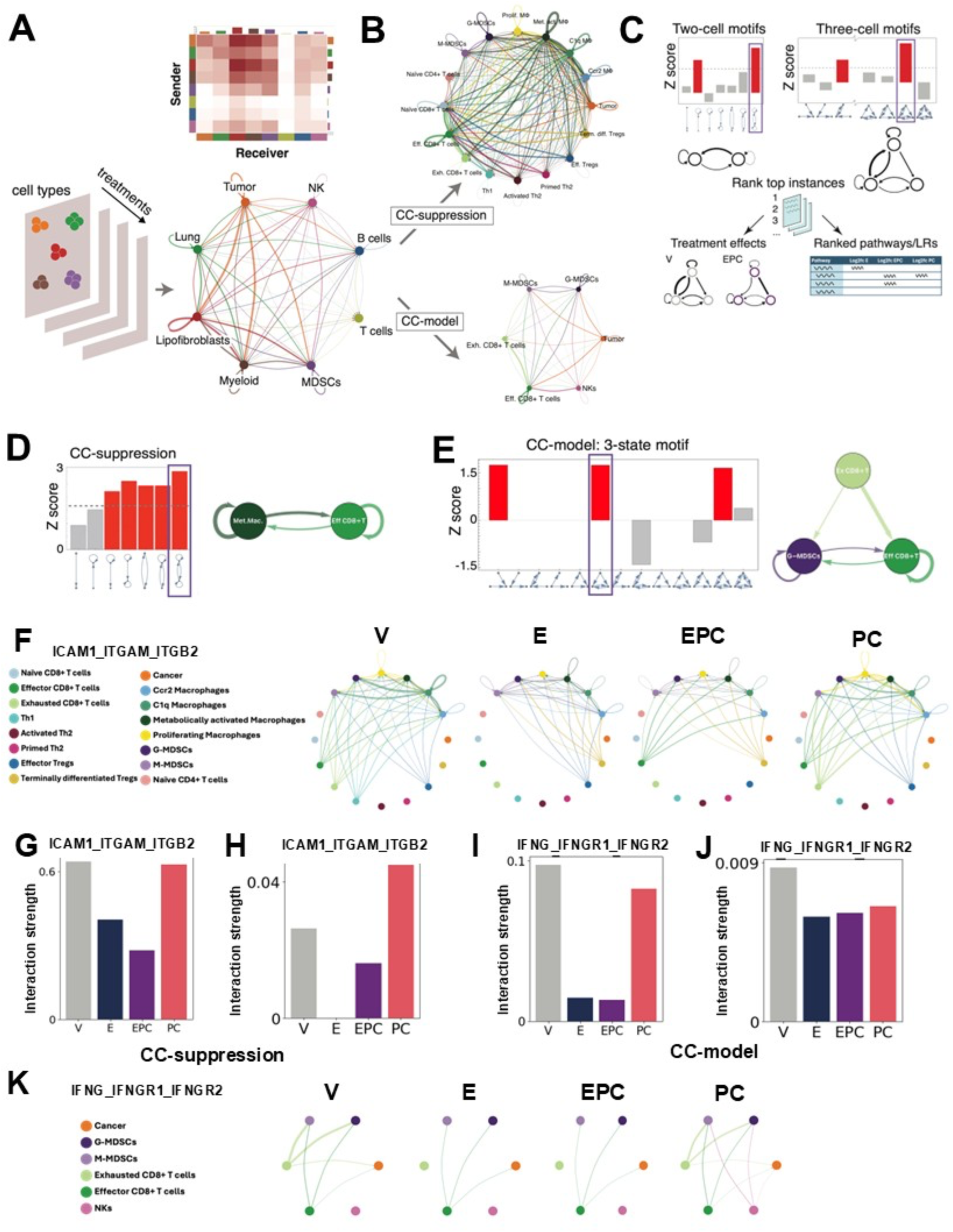
Cell-cell communication and cell circuits analysis of treatment effects in NT2.5-LM lung metastases. **A-C.** Schematic workflow of the cell-cell communication network analysis pipeline. **A.** High-level cell states were input to cell-cell communication network inference with CellChat, with heatmap and circos plots output. **B.** Two knowledge-guided analyses on subsets of cell states were performed: CC-suppression and CC-model. **C.** Overview of cell circuits pipeline. The input is a cell-cell communication network, and the outputs are top-ranked motifs and pathways/LR pairs. **D.** Top two-state motifs in CC-suppression, with the top instance of this motif being metabolically activated macrophages and effector CD8+ T cells (significant Z scores in red). **E.** The top three-state motifs from the CC-model involving G-MDSCs, effector CD8+ T cells, and exhausted CD8+ T cells. **F.** Circos plot displaying ICAM1_ITGAM_ITGB2 interactions between cells incorporated in the CC-suppression cell chat analysis with treatments. Arrows point from sources of *ICAM1* to cells interacting with *Icam1* via MAC-1 (*Itgam* + *Itgb2*). **G-H.** Bar plots showing the predicted interaction strength for ICAM1_ITGAM_ITGB2 between all cell populations within the CC-suppression comparison **(G)**, or between metabolically activated macrophages and effector CD8+ T cells **(H)** with treatments. **I.** Bar plots showing the predicted interaction strength for IFNG_IFNGR1_IFNGR2 between all cell populations within the CC-model comparison **(I)**, or between G-MDSCs, effector CD8+ T cells, and exhausted CD8+ T cells **(J)** with treatments. **K.** Circos plot displaying IFNG_IFNGR1_IFNGR2 interactions between cells incorporated in the CC-model analysis with treatments. Arrows point from sources of *Ifng* to cells interacting with *Ifng* via *Ifngr1* and *Ifngr2*. Increased arrow thickness indicates increases in interaction strength for panels **(F)** and **(K)**.

Given the complexity of cell signaling networks, even with reduced subsets of cell types (**Figure 4B**), we decomposed each based on an approach introduced in Mayer *et al*.^42^ that identified overrepresented motifs or “cell circuits” by studying all possible network wirings of two-node or three-node subgraphs of the directed, weighted cell-cell interaction graph (see Methods). Subgraphs that occurred more frequently than expected by chance were identified as cell circuits and ranked by their interaction strength (**Figure 4C**). Identification of two- or three-cell state circuits aids validation efforts, since functional studies are limited to the manipulation of only 2 (or max 3) cell types at once.

For CC-suppression, the top cell circuit for both 2-state and 3-state motifs was fully connected (i.e., bidirectional signaling between all cell types involved). Top-ranked instances of this motif by interaction strength revealed signaling between different myeloid and T cell subtypes, with effector CD8+ T cell---metabolically activated macrophage interactions at the top (**Figure 4D**), followed by C1q macrophages or G-MDSCs also interacting with effector CD8+ T cells (**Figure S10A**). For 3-state motifs, we found fully connected motifs consisting mostly of the same cell subtypes as for the 2-state motifs, the highest ranked consisting of metabolically activated and C1q macrophages interacting with effector CD8+ T cells (**Figure S10B**). The highest ranked motif, excluding cell subtypes from the same major types, was metabolically activated macrophages, G-MDSCs, and effector CD8+ T cells (**Figure S10C**).

In the case of CC-model using 2-state motifs, we found that the strongest motifs involved unidirectional signaling, with the strongest signaling occurring from NK cells to effector CD8+ T cells and from NK cells to G-MDSCs (**Figure S11A**). For the 3-state cell circuits in CC-model, several network motifs were statistically significant. We focused on the feedforward/feedback motif (**Figure 4E**), the top instances of which involved signaling between G-MDSCs, M-MDSCs, and effector CD8+ T cells (**Figure S11B**). Since differences between MDSC subtypes were unclear, for pathway analysis, we focused on the second-top motif, consisting of signaling between effector CD8+ T cells, exhausted CD8+ T cells, and G-MDSCs (**Figure 4E**).

### ICAM and IFN-II signaling between myeloid cells and T cells are significantly affected by combination treatment

We next identified which signaling pathways and ligand/receptor (L/R) pairs were commonly affected by E and EPC across multiple 2-cell circuits involving myeloid cells with CD8+ T cells. We looked at pathway changes involving metabolically activated macrophages and effector CD8+ T cells, G-MDSCs and effector CD8+ T cells, and macrophages and G-MDSCs (**Figure S11C-E**). The ICAM pathway was significantly altered by E and maintained by EPC treatment in all three of these circuits. Specific changes in L/R pairs that resulted in these pathway results are listed by treatment in **Supplementary File 3**. Given numerous significantly differentially regulated L/R pairs affected by treatment, we focused on biologically relevant pairs that have been reported to modulate the immune response to cancer. ICAM1_ITGAM_ITGB2 interactions decreased with E and EPC treatments overall in CC-suppression, and specifically between total macrophages and effector CD8+ T cells (**Figure 4F-H**). IFNG_IFNGR1_IFNGR2 interactions decreased with E and EPC treatments across all cell types within the CC-model and specifically between G-MDSCs and exhausted CD8+ T cells (**Figure 4I-K**). The active form of MAC-1 (*Itgb2_Itgam*) expressed on macrophages inhibits antigen presentation^43^, while IFN-ɣ maintains MDSC suppression of T cells and induces the production of immunosuppressive iNOS in MDSCs^44^. These findings suggest that EPC increased antigen presentation by macrophages and decreased suppression of T cells by myeloid cells.

### ICAM-1 and IFN-ɣ blockade interrupt myeloid---CD8+ T interactions and cause functional changes affecting CD8+ T cell proliferation

To investigate the functional effects of EPC treatment on different myeloid cell populations identified within the lung metastatic TME of NT2.5-LM and 4T1 mice, we conducted *ex-vivo* co-cultures with CD8+ T cells. While G-MDSCs inhibited T cell proliferation in both models (**Figure 5A, S12A**), total macrophages only suppressed T cell proliferation when obtained from 4T1 mice (**Figure 5B**). Given the known plasticity of macrophage populations^19^, we investigated changes in expression of MHC-II on macrophages when processed and analyzed immediately (15 hours) compared to the following day (38 hours). MHC-II expression increased significantly over time, which could contribute to the observed variability in CD8+ T cell suppression by macrophages (**Figure S12B**). Monocytes and neutrophils did not affect T cell proliferation (**Figure 5C**). *In-vivo* treatment of mice followed by *ex-vivo* co-cultures did not affect G-MDSC or macrophage inhibition of CD8+ T cell proliferation in either model (**Figure 5D-E**). These findings suggest that treatment with E and EPC does not rescue T cell proliferation within lung metastases, unlike what we previously observed with myeloid cells isolated from treated primary tumors^10^. To validate the predicted functional effects of EPC on ligand-receptor pairs identified in our scRNA-seq CellChat analysis, we blocked ICAM-1 and IFN-ɣ receptor *in ex-*vivo co-culture assays. Inhibition of ICAM- 1 on T cells stimulated their proliferation when co-cultured with total macrophages from NT2.5- LM metastases, but not those from 4T1 (**Figure 5F**). In addition, expression of PD-L1 decreased in macrophages following ICAM-1 blockade, concordant with functional changes in the NT2.5-LM model (**Figure S12C**). However, ICAM-1 blockade *in-vivo* did not affect survival with or without ICIs, as opposed to with EPC (**Figure S12D**). These results suggest that the entinostat-driven decrease in ICAM-1 signaling promotes a functional shift from pro-tumor to pro-inflammatory macrophages, which has no survival benefit on its own. Inhibition of IFN-ɣ signaling rescued CD8+ T cell proliferation when co-cultured with G-MDSCs in both models (**Figure 5G**), suggesting that decreased IFN-ɣ production observed with EPC could contribute to the improved response to checkpoint inhibition.

**Figure 5.**
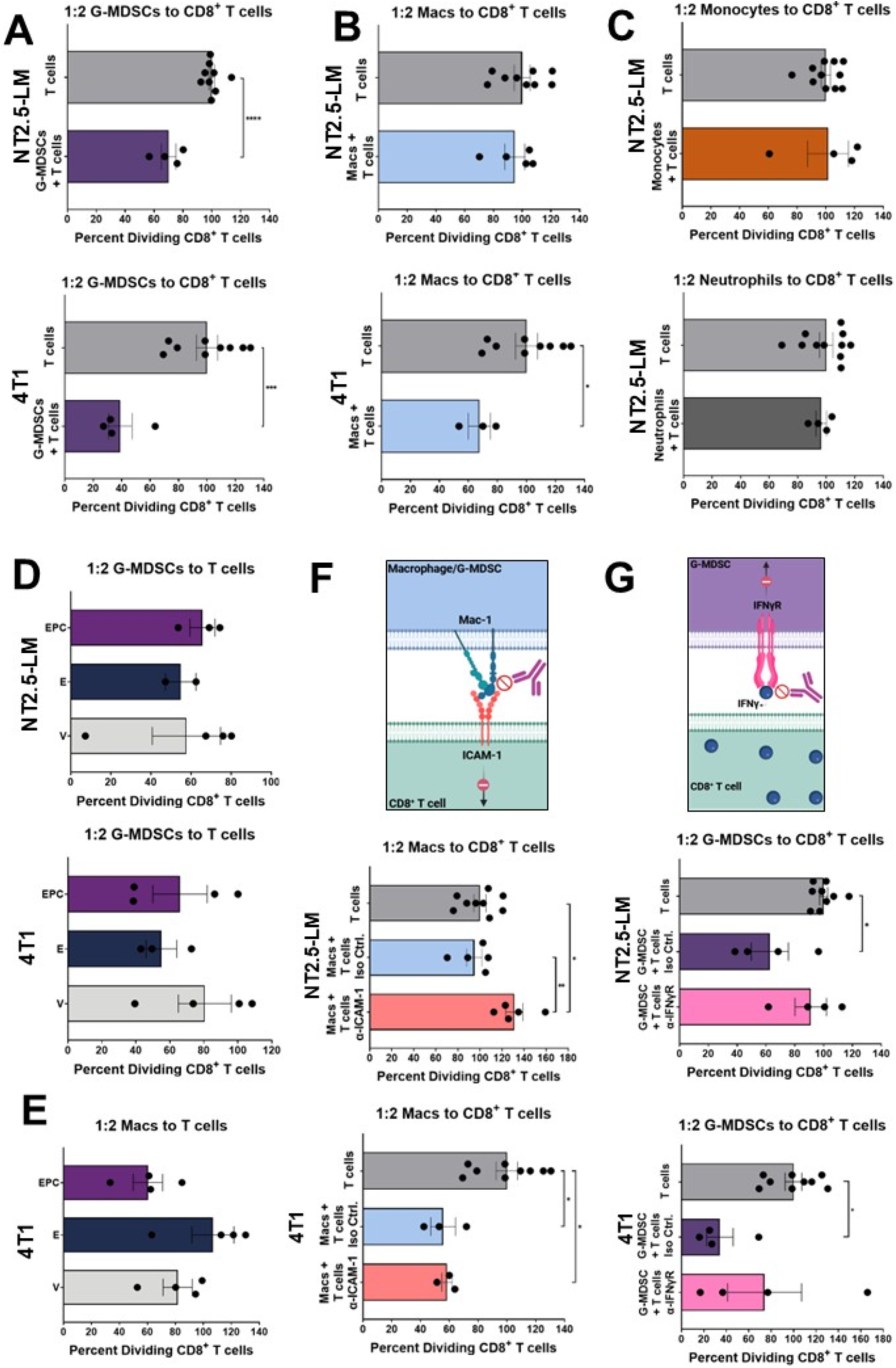
CD8+ T cell co-culture assays to investigate changes in the function of suppressive myeloid cells with treatment. **A-B.** Bar graphs depicting the proportion of CD8+ T cells undergoing proliferation, either cultured alone (T cells) or in co-culture with G-MDSCs (**A**; n=4/group) or total macrophages (**B**; n=3-4/group ) from NT2.5-LM (top) or 4T1 (bottom) lung metastases. **C.** Bar graphs depicting the proportion of CD8+ T cells undergoing proliferation in co-culture with monocytes (top) or neutrophils (bottom) from NT2.5-LM lung metastases (n=4/group). **D.** Proportion of CD8+ T cells undergoing proliferation when in co-culture assays with G-MDSCs obtained from NT2.5-LM (top) or 4T1 (bottom) lung metastases from mice treated with vehicle (V), entinostat (E), or entinostat + anti-PD-1 (P) + anti-CTLA-4 (C). n=2-4/group. **E.** Proportion of CD8+ T cells undergoing proliferation when in co-culture assays with total macrophages obtained from 4T1 lung metastases from mice treated with V, E, or EPC (n=4/group). **F.** Depiction of blockade of ICAM-1 signaling in macrophages using neutralizing antibodies targeting ICAM-1 (top) and proportion of CD8+ T cells undergoing proliferation when in co-culture with total macrophages from NT2.5-LM (middle) or 4T1 (bottom) lung metastases. Mac-1 = CD11b (*Itgam*) + Itgb2. **G.** Depiction of blockade of IFNɣ signaling in G-MDSCs using neutralizing antibodies targeting IFNɣ receptor (IFNɣR; top) and proportion of CD8+ T cells undergoing proliferation when in co-culture with G-MDSCs from NT2.5-LM (middle) or 4T1 (bottom) lung metastases. Statistically significant differences were determined using the unpaired t-test for panels A-C and the one-way ANOVA for panels D-G. *p<0.05, **p<0.01, ***p<0.001. Antibody panels used for flow cytometry are found in Supplementary File 2. CD8+ T cell proliferation in each group was normalized to the T cells alone control for all experiments.

### Mathematical modeling of TME dynamics highlights that dual modulation of CD8+ T cell activation and inhibition by MDSCs are necessary for tumor shrinkage

Previously, we developed a tumor--immune model to study the impact of MDSCs on TME dynamics in the presence of CD8+ T cells and NK cells and found two parameters, MDSC recruitment to the TME and NK cell suppression by MDSCs, to be critical for new metastatic growth^41^. We modified our previous model to encapsulate the metastatic TME by adjusting parameters and initial conditions to fit the context of established lung metastases undergoing treatment (see Methods), after confirming that the cell types accounted for in the model were present in our scRNA-seq dataset with numerous pathways mediating cell-cell communication between cell types (**Figure 4A**). The previous model was changed such that there was no MDSC recruitment delay to the tumor microenvironment (𝜏 = 0 ), tumor cell population was at steady state, and the immune cell populations were initialized according to cell composition determined via scRNA-seq in the untreated TME.

To test how tumors respond to treatment effects mediated by specific cell-cell interactions and changes predicted by our cell circuits analysis above, we identified and scaled cell-cell interaction parameters in the model according to cell-cell communication interaction scores^31^. Analysis of the relevant cell circuits and associated LR pairs/pathways determined two dominant effects of EPC treatment: decreased MDSC inhibition of CD8+ T cells (β_4_ → β_4_ × 0.38) and increased activation of CD8+ T cells (α_6_ → α_6_ × 1.4) (**Figure 6A-B**).

**Figure 6.**
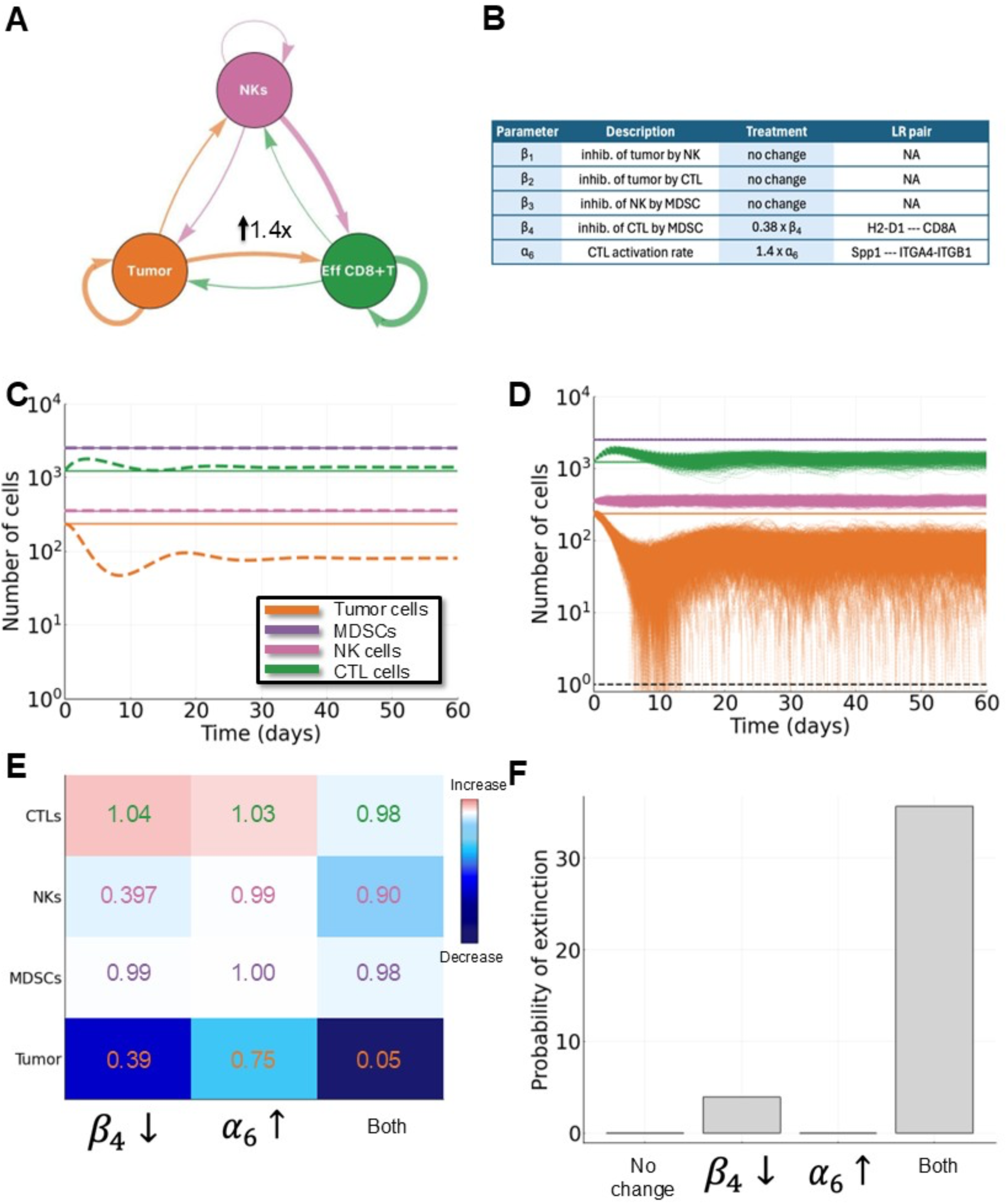
Mathematical modeling predicts a decrease in cancer cells with treatment. **A.** Schematic of a cell circuit comprising cancer cells, NK cells, and effector CD8+ T cells. Comparing this cell circuit between V and EPC treatments suggests that alpha_6 (CTL stimulation by tumor-NK interaction) increases with treatment. **B.** Table of important model parameters and how they change with treatment. **C.** Deterministic simulation of the ordinary differential equation mathematical model with (dashed lines) and without (solid lines) treatment. The tumor population decreases with treatment. **D.** 10^3 simulations of the stochastic differential equation mathematical model. **E.** Heatmap showing overall differences in the four cell populations included in the model under different treatment effects (parameter changes). **F.** Bar plot showing the probability of extinction for different treatment effects (parameter changes). Tumors are declared extinct when their population size is below one cell.

When both these treatment effects were modeled simultaneously, we observed a large tumor response to treatment (**Figure 6C-D**), where, in the stochastic case (**Figure 6D**), some tumors go extinct (defined as <1 tumor cell remaining). Comparison of individual vs. dual effects (**Figure 6E-F**) found that only with the dual effect was the tumor size dramatically reduced after treatment compared to baseline (95% decrease; deterministic case). In the stochastic case, this led to a probability of tumor extinction of 35.7% (**Figure 6F**), where the best that can be achieved when only modulating a single parameter is 3.9%.

### Analysis of patient biopsies via spatial proteomics validates predicted myeloid and B cell phenotypes with EPC treatment in clinical responders

We turned to IMC biopsies from our clinical trial (NCI-9844) to examine changes in immune cell infiltration and interactions with treatment. Using a panel of 34 antibodies for epithelial cells, stromal cells, immune cells, and functional markers (**Figure S13B**), we aggregated cell populations with similar patterns of protein expression into cell categories as described in the literature^45–51^, with each immune cell population showing characteristic expression levels of at least one signature cell marker (**Figure S13C**, underlined) except pro-tumor macrophages, which represent a heterogeneous population of subclusters. The tissue biopsy site influenced immune heterogeneity, as exemplified by large DC proportions in breast tissues (**Figure S13A**). An analysis of high-level immune cell types revealed no differences in proportions between responders and non-responders at any time point (**Figure 7A**). To compensate for small sample sizes, we pooled samples across time points, resulting in increased proportions of naive CD4+ T cells and decreased proportions of pro-tumor and resting macrophages in responders compared to non-responders, although the decrease in naïve CD4+ T cells and pro-tumor macrophages did not meet significance criteria (p<0.0841; **Figure 7B**). Stratifying the data by treatment produced similar trends at all time points (**Figure S14A**). To determine the effects of EPC on immune cells associated with treatment response, we analyzed changes in immune cell proportions at week 8 compared to baseline. EPC treatment increased the proportions of total CD8+ T cells in responders specifically and mature DCs in all samples, although the increase in CD8+ T cells did not meet significance criteria (p=0.0698; **Figure 7C-D**).

**Figure 7.**
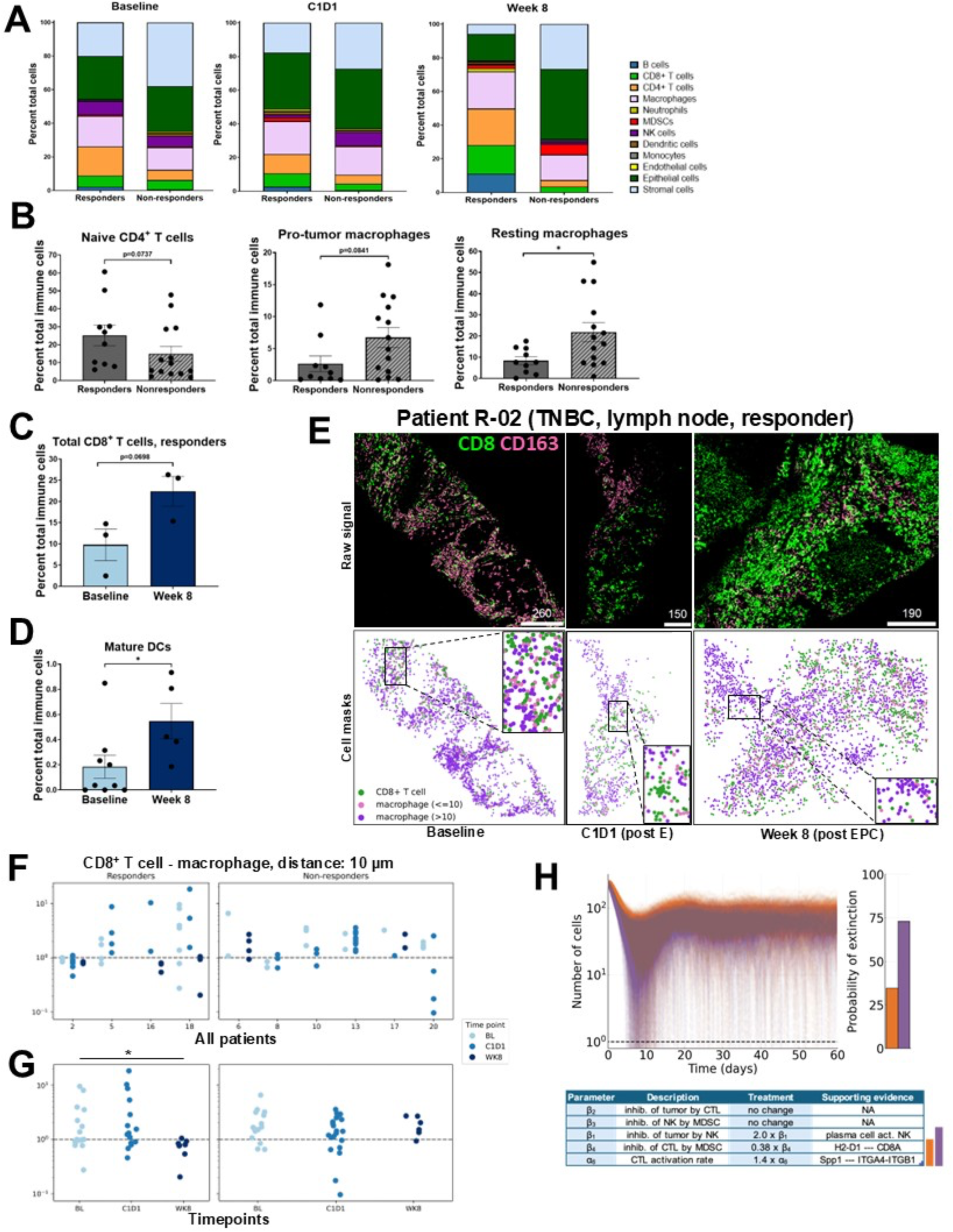
Spatial proteomics identifies changes in immune cells in patient tumors with EPC treatment. **A.** Proportions of major cell populations identified via imaging mass cytometry (IMC) in responders (n=4) versus non-responders (n=6) at baseline, post-entinostat (C1D1) and at week 8 (following the addition of anti-PD-1 + anti-CTLA-4 treatments), excluding cells that did not express any cell markers included in the IMC panel, classified as “other”. Patient tumor tissues representing breast tumors and metastatic tissues from lungs, liver, chest wall, and lymph nodes were included in the analysis. **B.** Proportions of naïve CD4+ T cells, pro-tumor macrophages, and resting macrophages in responders (n=10) versus non-responders (n=14) when analyzing samples from all timepoints. *p<0.05, as determined via the Mann-Whitney test. **C.** Proportions of total CD8+ T cells in responders at baseline compared to week 8 (n=3/group). The unpaired t-test was used to determine statistically significant differences between groups. **D.** Proportions of mature dendritic cells (DCs) in all samples, regardless of response, at baseline (n=9) compared to week 8 (n=5). Statistically significant differences were determined via the Mann-Whitney test. **E.** Top: Visualization of raw signals for CD8 (green) and CD163 (pink) in IMC images from patient R-02’s lymph node samples at baseline, C1D1, and week 8. Bottom: Visualization of IMC masks on the tissues shown above, demonstrating the locations of CD8+ T cells (green dots) and macrophages at distances above 10 µm (purple) and below 10 µm (pink). **F-G.** Co-occurrence scores between CD8+ T cells and total macrophages for individual patients **(F)** and co-occurrence scores between CD8+ T cells and total macrophages for all responders (7-14 ROIs/group) and all non-responders (6-19 ROIs/group; **G**). The Kruskal-Wallis test determined significant differences in co-occurrence scores between timepoints. **H.** 10^3 simulations of the stochastic differential equation mathematical model for orange lines: treatment effects of only alpha_6 (CTL stimulation by tumor-NK interaction) and beta_4 (inhibition of CTL by MDSCs), and purple lines: treatment effects of alpha_6, beta_4, and beta_1 (inhibition of tumor by NK). The change in beta_1 for patient R-2 is supported by the dramatic increase in plasma cells, which activate NK cells against tumor cells. The bar plot shows a dramatic increase in the probability of extinction for simulated tumors with increased beta_1 parameter.

Response to checkpoint inhibition has also been associated with changes in physical cell-cell interactions^52,53^. Via co-occurrence analysis, we observed increased distances between CD8+ T cells and total macrophages at week 8 compared to BL (**Figure 7E-F**). Analysis over time revealed increased distances (decreased co-occurrence) between CD8+ T cells and total macrophages in responders at week 8 compared to baseline (**Figure 7G**). In support of these findings, responders demonstrated increased expression of CD137 in CD8+ T cells and granzyme B in NK cells, both markers of activation^54^ (**Figure S14B-C)**. HLA-DR expression was higher in pro-inflammatory macrophages at baseline and week 8 in responders compared to non-responders (**Figure S14D**). Also, responders had decreased expression of PD-L1 in Tregs at week 8 versus baseline (**Figure S14E**).

The trending increase in plasma cell proportions with treatments observed in responders (**Figure** S7F) was recapitulated in patient R-02, who displayed an increase in plasma cell proportions with E and EPC treatment (**Figure S13A**). To assess the impact of this treatment effect, we simulated our mathematical model in a new parameter regime: combining the effects of EPC treatment from our analysis above (**Figure 6**) with the predicted impact of an increase in plasma cells on cytotoxic NK cell activation. Over a 60-day window, we observed an increase in the probability that a tumor will go extinct (purple vs orange bar, **Figure 7H**). This highlights the potential treatment benefit of increased plasma cell proportions observed in responders and is in agreement with the response observed in patient R-02 following treatment.

## DISCUSSION

Checkpoint inhibitors successfully induce lasting anti-tumor immunity when they increase infiltration and activation of CD8+ T cells, which can lead to response in many cancer types, including metastatic breast cancer^55^. Entinostat has been shown to decrease myeloid suppression through effects on MDSCs, contributing to an enhanced CD8+ T cell response^8,10,12^. However, this does not fully account for the observed complexity of sensitization of the metastatic TME to checkpoint inhibition. Our knowledge-guided categorization of immune cell populations within breast-to-lung metastases from two murine models analyzed by scRNAseq and flow cytometry revealed significant shifts in subpopulations of immune cells with EPC treatment, including CD8+ T cells, Th2 cells, and B cells. While the changes observed in activated Th2 proportions were caused by entinostat alone and sustained following the addition of dual ICIs, the effects of EPC on CD8+ T cells and B cells were produced via a synergistic effect between E and PC treatment. Cell circuits analysis identified the ICAM and IFN-II pathways as downregulated by combination treatment, and we found that interruption of ICAM-1 and IFN-ɣ signaling in co-culture assays involving CD8+ T cells and macrophages/G-MDSCs increased T cell proliferation. In addition, the decrease in physical interactions between macrophages and CD8+ T cells observed in patient tumors with EPC in responders implies that these interactions could be a crucial target for an effective anti-tumor response with ICIs. Mathematical modeling of TME dynamics confirmed that increased CD8+ T cell activation by B cells and checkpoint inhibition and decreased MDSC inhibition of T cells are both necessary for tumor shrinkage, highlighting the necessity for drugs that can target multiple cell types or molecules to achieve an anti-tumor immune response. Analysis of patient biopsies via imaging mass cytometry confirmed an increase in plasma cell generation and B cell activation markers with EPC. We also observed altered immunosuppression, as the distances from CD8+ T cells to myeloid cells increased in responders, as predicted by preclinical studies.

Traditionally, studies into mechanisms of response seek to identify individual genes, pathways, protein targets/receptors, or cell types necessary or sufficient to drive the observed outcome. Our study reveals intrinsic limitations with this approach. In the case of immunosuppressed tumors where a nonspecific drug such as the epigenetic modulator entinostat has an effect, there are many co-occurrent sources of intrinsic resistance that need to be addressed^4–7^. We observed increased activation of Th2 cells with EPC in both NT2.5-LM and 4T1 models, but increased effector CD8+ T cell proportions only in the 4T1 model, which may be more responsive to EPC treatment^8^. We observed drastically reduced IFN-ɣ signaling via CellChat in NT2.5-LM lung metastases via scRNA-seq and a Th2-driven response elicited by EPC in both preclinical models. This could however be due to background: both mouse models are from BALB/c, which are known to display predominantly Th2-driven responses^56,57^. Further testing of patient samples and mouse studies using the 4T1 model will serve to verify if the decrease in IFN-ɣ production observed with EPC is specific to the NT2.5-LM model or applies generally.

Treatment-induced changes in B cell function and immunoglobulin production with EPC in both mouse and patient analysis suggested a favorable prognosis and a positive response to ICIs^58^. The higher MHC-II expression in B cells from NT2.5-LM tumors and the higher B cell CD137 expression via spatial proteomics on patient biopsies suggested an upregulation of antigen presentation from B cells to T cells with EPC. This is in line with a cervical cancer model, where B cells triggered anti-tumor responses better than dendritic cells in an MHC-II-dependent manner^59^. Furthermore, we found evidence that EPC drives an antibody class switching to IgG1, likely via stimulation of a Th2 response, which could lead to the elimination via NK cells and macrophages expressing IgG-specific FcR receptors^60^. In patient biopsies, increased proportions of B cells were observed in responders compared to non-responders, suggesting that an intrinsically higher B cell tumor-infiltration rate could facilitate EPC efficacy. Together, the effects observed with the combination of entinostat and checkpoint inhibitors suggests that effective treatment response requires a multi-pronged approach, at once decreasing immunosuppression and stimulating effector T cells/B cells in the TME.

A cancer systems immunology approach was crucial to identify mechanisms of response to combination therapy. The complexity of the TME is vast: with 39 cell states, there exist 9,139 three-state signaling motifs. Through cell-cell communication network and cell circuit inference, we were able to identify a small number of top motifs associated with treatment. Due to the rarity of B cells in the lung microenvironment, quantifying differences in numbers or transcriptional states would not be possible without sequencing ∼10^5 cells. By integrating mathematical models with data-driven insights (a growing subfield of mathematical oncology^61–63^) it was possible to predict quantitative tumor responses to specific combinations of TME parameter perturbations, especially those controlling immune suppression. In combination, our preclinical and mathematical results were invaluable in guiding the analysis of IMC patient biopsies.

In conclusion, we have described how mechanisms of immune suppression mediated primarily by macrophages and MDSCs on T cells and B cells can be overcome with combination therapy via checkpoint inhibitors combined with entinostat. This was made possible only by taking a cancer systems immunology approach involving the integration of scRNA-seq, cell-cell communication analyses, mathematical modeling, functional immunology, and follow-up validation studies on patient biopsies. Beyond the impact of the current mechanisms elucidated, this study serves as a framework for other investigations into mechanisms of response to new therapeutics in complex TMEs.

## METHODS

### Mice

Female NeuN mice (Jackson Labs strain #002376) were used for studies with the NT2.5-LM cell line, which were bred in-house via brother-sister mating in the animal housing facility at the University of Southern California. Female BALB/c mice used for studies with the 4T1 cell line were purchased from Jackson Labs (strain #000651). All mice were housed under pathogen-free conditions and handled according to protocol 21770 approved by the USC Institutional Animal Care and Use Committee, following guidelines by the American Association of Laboratory Animal Committee policies.

### Cell lines

NT2.5-LM is a rat HER2/neu+ breast cancer cell line derived from NT2.5 that spontaneously metastasizes to the lungs of NeuN mice with high penetrance^14^. 4T1 is a triple-negative breast cancer cell line that also shows high rates of spontaneous lung metastasis^8^. Both NT2.5-LM and 4T1 cells were cultured in RPMI-1640 base media (Gibco, cat. # 11875-093) supplemented with 20% fetal bovine serum (Gemini, cat. # 100-106), 1.2% HEPES (Gibco, cat. # 15630-080), 1% l- glutamine (Gibco, cat. # 25030-081), 1% non-essential amino acids (Gibco, cat. % 11140-050), 1% sodium pyruvate (Sigma, cat. # S8636), 0.5% penicillin/streptomycin (Gibco, cat. # 15140- 122), and 0.2% insulin (NovoLog, cat. # NDC 0169-7501-11) at 37 °C and 5% carbon dioxide. Cell media used for co-culture assays with CD8+ T cells was composed of RPMI-1640 base media supplemented with 10% fetal bovine serum, 1.5% HEPES, 1% l-glutamine, 1% non-essential amino acids, 1% sodium pyruvate, 1% penicillin/streptomycin, and 0.0004% beta-mercaptoethanol (Sigma, cat. # M3148). Cell lines were tested for mycoplasma contamination yearly. All cells were thawed and kept in culture for a minimum of five days before injection into mice or for *in vitro* experiments.

### *In-vivo* Experiments

For spontaneous metastasis studies described in Figures 1C-H, 2A-G, S2A-D, S3A-G, S6A-D, S6F-J, 3A-C, 3F-I, S7A-E, S8-S11, 4A-K, 5I-J (4T1), and 6A-F, eight-to-twelve week-old female NeuN mice or BALB/c mice were injected with 100,000 NT2.5-LM cells (NeuN) or 50,000 4T1 cells (BALB/c) in the 4^th^ mammary fat pad. For experimental metastasis studies described in Figures 1A, 2H, S2E, S4, S5, S6E, 3D-E, 3J, S12, and 5A-K, eight-to-twelve week-old female NeuN mice or BALB/c mice were injected with 100,000 NT2.5-LM cells (NeuN) or 4T1 cells (BALB/c) in the tail vein. For studies involving treatments with entinostat (Syndax Pharmaceuticals) and ICIs, mice were administered 5 mg/kg entinostat five times/week via oral gavage and 100 µg/dose of anti-CTLA-4 (BioXCell cat. # BE0131) and anti-PD-1 (BioXCell cat. # BE0146) twice a week via intraperitoneal (IP) injection. For studies involving anti-ICAM-1 treatment, anti-ICAM-1 was administered at a dose of 100 µg per injection, five times/week via IP injection. Entinostat treatment was started one day before ICIs therapy, starting on day 21 post-injection for the survival experiment in Figure 1A, day 24 post-injection for the survival experiment in Figure 5K, day 17 post-injection for the scRNA-seq experiment with NT2.5-LM, day 10 post-injection for all co-culture assays with NT2.5-LM described in Figure 5 and S12, day 2 post-injection for the flow cytometry experiment in 4T1, and day 36 post-injection for the co-culture assays in 4T1 with neutralizing antibody treatments (Figure 5I-J). Treatments were maintained for a minimum of three weeks for all *in-vivo* experiments. For survival studies, mice were observed daily for symptoms of lung metastasis, including labored breathing, ruffled coat, and reduced activity, and euthanized once they presented clear symptoms. Mice were considered to reach the endpoint for survival if lung metastases were detectable by visual inspection during dissection or pleural effusion was observed, which was detected in most cases of lung metastasis. For survival studies using a spontaneous model of lung metastasis, tumors were resected from mice on day 12 post-injection for NT2.5-LM or days 29-30 post-injection for 4T1. For studies involving flow cytometry analysis of lung metastases or co-culture assays with CD8+ T cells, mice were euthanized 1 day before starting to succumb to disease based on previous studies, lungs were inflated with sterile RPMI-1640 media, dissected, and stored on ice submerged in media during processing of samples until analysis or 37 °C incubation started for co-culture assays. For the opsonization experiments described in Figure S3J and S7E, blood was collected from mice via cardiac puncture following euthanasia. Blood samples were incubated at room temperature for 30 minutes, centrifuged at 1500 g for 10 minutes, transferred to microcentrifuge tubes, and stored at –80 °C.

### Processing of lung metastases from mice for flow cytometry and MACS

Lung metastases from NeuN (NT2.5-LM) or BALB/c mice (4T1) were dissected from lungs using fine-tipped scissors under sterile conditions inside a biosafety level 2 cabinet and digested using the Miltenyi mouse tumor dissociation kit (Miltenyi, cat. # 130-096-730) in combination with the Miltenyi Octo dissociator, following the manufacturer’s directions. Samples were processed through a 70 µm strainer into single-cell suspensions, and red blood cells were lysed via a five-minute incubation in ACK lysis buffer (Gibco, cat. # A1049201) before adding antibodies for flow cytometry or magnetic-activated cell sorting (MACS).

Samples composed of lung lobes all containing lung metastases described in Figure S12B-C were split into 2 and either analyzed immediately following lung dissection or stored overnight at 4 °C by submersion in MACS® Tissue Storage Solution (Miltenyi, cat. # 130-100-008) before analysis via flow cytometry.

### Flow cytometry, FACS, and MACS

Following the processing of NT2.5-LM or 4T1 lung metastases into single-cell suspensions and lysis of red blood cells, samples to be analyzed by flow cytometry or samples to be FACS-sorted for co-culture assays were transferred to u-bottom 96-well plates at a minimum of 200,000 cells/well and a maximum of 1,000,000 cells/well, stained using the LIVE/DEAD Fixable Aqua Dead Cell Stain kit (ThermoFisher Scientific, cat. # L34966) by resuspending cells in 100 µL dye at 1:1000 dilution in FACS buffer (10% FBS in PBS) and incubating at 4 °C, protected from light exposure, for 30 minutes. Samples were blocked with 10% FBS during dead cell staining and during washes, and Fc receptors were blocked via incubation of samples with 1:50 anti-mouse CD16/CD32 (Biolegend, cat. # 101302) at room temperature for 10 minutes. Samples were stained for one hour at 4 °C with a panel of antibodies targeting proteins specific to different immune cell types or isotype control antibodies (all antibody panels used for flow cytometry including manufacturer and catalogue numbers for each target and isotype antibody are found in S**upplementary File 2),** as identified in the literature (**Supplementary File 1;** includes references) and in immune cell populations via our scRNA-seq analysis. For intracellular staining, samples were fixed and permeabilized using the Foxp3/Transcription Factor Staining Buffer Set (eBioscience™, cat. #00-5523-00) or the Cyto-Fast™ Fix/Perm buffer set (Biolegend, cat. #426803) following the manufacturer’s directions prior to a 10-minute FcR block step at room temperature and a one-hour antibody staining step at 4 °C. Samples were protected from light exposure and kept on ice, when possible, throughout staining and up until the moment of analysis.

Samples analyzed via MACS were incubated with magnetic bead-conjugated antibodies included in the mouse Myeloid-Derived Suppressor Cell Isolation Kit (Miltenyi, cat. # 130-094-538), and G-MDSCs were isolated following the manufacturer’s directions.

### Opsonization assays

For the experiment described in Figure S7E, NT2.5-LM cells were thawed and cultured for 1 week prior to staining dead cells and conducting blocking steps as described for flow cytometry experiments (above). NT2.5-LM cells were incubated for one hour at 4 °C with serum diluted 1:100 in FACS buffer from healthy mice or mice harboring NT2.5-LM lung metastases treated with V, E, or EPC. A secondary PE-conjugated anti-mouse IgG (Invitrogen, cat. # 12-4010-82) was used to determine NT2.5-LM-targeting IgG concentrations in serum by quantifying the median fluorescence intensity (MFI) signal for PE in NT2.5-LM cells gated as CD45-HER2+ single, live cells using flow cytometry.

For the experiment described in Figure S3J, the level of endogenous IgG opsonization of HER2+ cancer cells in NT2.5-LM lung metastases was analyzed for differences with V, E, PC, and EPC treatments. Ten frozen NT2.5-LM lung metastasis samples from an experimental lung metastasis study and 14 frozen samples from a spontaneous lung metastasis study (n=2-5/treatment group) were thawed, dead cells were stained, and blocking steps were conducted as described for flow cytometry experiments (above). Cancer cells within NT2.5-LM lung metastases were gated as CD45-HER2+ single, live cells, and PE-conjugated anti-mouse IgG was used to detect levels of endogenously bound IgG via quantification of PE MFI signal.

### *Ex-vivo* CD8+ T cell co-culture assays for proliferation

For all co-culture experiments described in Figure 5, myeloid cells were obtained via FACS-sorting of single cell suspensions of NT2.5-LM lung metastases and co-cultured *ex-vivo* with splenic CD8+ T cells obtained from healthy mice. CD8+ T cells were isolated from spleens using the mouse CD8a+ T Cell Isolation Kit (Miltenyi, cat. # 130-104-075) following the manufacturer’s directions, stained with CellTrace™ CFSE dye (ThermoFisher Scientific, cat. # C34554), and stimulated with mouse T-Activator CD3/CD28 Dynabeads™ (ThermoFisher Scientific, cat. # 11452D). NT2.5-LM lung metastases were prepared for FACS sorting as described under the flow cytometry protocol (above), then G-MDSCs were sorted as CD45+CD11b+F4/80-Ly6G+Ly6C+ and total macrophages were sorted as CD45+CD11b+F4/80+Ly6G-MHCII+FLT3- single, live cells. Monocytes were gated as CD45+CD11b+F4/80+Ly6G-MHCII-FLT3- and neutrophils were sorted as CD45+CD11b+F4/80+Ly6G+ single, live cells. CD8+ T cells were analyzed via flow cytometry by gating for CD3+CD11b- or CD8+CD11b- single, live cells, and cell proliferation was determined via quantification of median MFI for CFSE dye after 60-72 hours of incubation at 37 °C. All co-cultures were set up at ratios of 1:2 CD8+ T cells to target cells except for the G-MDSC co-cultures illustrated in Figures S12A and S12C, which were set up at 1:1, 1:2, 1:4, and 1:8 ratios using a sterile 96-well flat-bottom, non-tissue culture-treated plate. A minimum of 10,000 and a maximum of 20,000 CD8+ T cells were plated per well for each co-culture assay. Samples of lung metastases and FACS-sorted lipofibroblasts, monocytes, neutrophils, G-MDSCs, or macrophages from NT2.5-LM and 4T1 lung metastases, along with splenic CD8+ T cells, were kept on ice until the co-culture assays were set up and placed in the incubator at 37 °C.

### Spatial Proteomics

Sections from formalin-fixed, paraffin-embedded patient tumor tissues from NCI-9844 representing primary breast tumors and metastatic tissues from chest wall, liver, lung, and lymph node from eleven patients with either estrogen receptor-positive or triple-negative breast cancer were analyzed via IMC using the 5 general markers and 34 metal-isotope tagged antibodies listed in **Figure S13B** for epithelial cells, stromal cells, immune cells, and functional markers as described by Barbetta *et al*., 2024^64^. The general markers included two iridium nuclear stains, 2 2 ruthenium tissue stains, and the PM4 tissue stain included in the Maxpar IMC Cell Segmentation Kit (TIS-00001). All tissues, including a total of 114 ROIs, were checked by a pathologist to ensure that only samples containing tumor tissue would be incorporated into the analysis. Sample details are outlined in **Figure S13A**. Pre-processing of raw IMC images for channels spillover, denoising, and aggregates was performed using the IMmuneCite-IMClean pipeline. Mesmer (DeepCell) was used to perform cell segmentation after running samples through the Bodenmiller Steinbock pipeline^65^. Image preprocessing was performed using Python (version 3.12.10).

A total 530,287 cells were identified in all samples. Cells with similar patterns of protein expression were aggregated into high-order clusters, from which subclusters representing individual cell populations were derived. Subclusters were binned into cell categories based on canonical marker expression to define immune cell types described in the literature^45–51,66–78^, with each subcluster showing characteristic levels of at least one signature cell marker for immune cells (**Figure S13C**, underlined), except early MDSCs^79^, for which there is no established cell marker and were classified as CD33+CD14-CD15-, and pro-tumor macrophages, which represent a heterogeneous population of subclusters each expressing different combinations of markers associated with a pro-tumor phenotype in macrophages, with the majority expressing CD163. Subclustering for each high-order cluster was performed using the FlowSOM algorithm^80^ implemented by the CATALYST package in R^81^. Expression data of cells in one high-order cluster was provided to each run of FlowSOM, along with a list of markers for potential subclusters. Nine subclusters were obtained for each high-order cluster and were annotated with subcluster labels based on their expression profile.

### Determination of TLS structures in Spatial Proteomics

High-order clusters were visualized on tissue masks for baseline, C1D1, and week 8 timepoints for patients that had detectable germinal centers or tertiary lymphoid structures (TLS): R-06, R- 08, R-10, R-02, and R-16. Samples were not obtained for patients R-08 and R-10 at week 8 and patient R-16 at baseline. Cells that did not fall under any of the categories listed due to a lack of cell marker expression were pooled into a group labeled “other.” A TLS was defined as an aggregation of 8 or more B cells/plasma cells within a 125 µm radius in non-lymph node tissues.

### Differential expression analysis of marker expression in spatial proteomic data

For each marker and each subset of cells (either all cells or a specific cluster), t-tests were performed to assess differences in marker expression. The t-tests compared differential expression between time points (baseline vs C1D1, baseline vs week 8, C1D1 vs week 8) within responders or non-responders, as well as between responders and non-responders at each time point. P-values from each comparison, for the same marker and cell subset, were adjusted using the Bonferroni correction.

### Co-occurrence analysis for high-order clusters in spatial proteomics

Co-occurrence analysis was performed on all high-order clusters from the IMC data using the SquidPy package^82^. For each IMC sample (ROI), one co-occurrence score was calculated between each pair of high-order clusters A and B by *P*(*B*|*A*)/*P*(*B*) , where *P*(*B*|*A*) is the conditional probability of observing cells from B within 10 µm of any cell from A, and *P*(*B*) is the general probability of observing cells from B within any circle of radius 10 µm. A higher co-occurrence score between A and B indicates a higher proximity of cells from the two high-order clusters. The Kruskal-Wallis test was performed on scores between CD8+ T cells and macrophages among all time points (baseline, C1D1, week 8), for either all responders or all non-responders, with the null hypothesis that scores from all the time points have equal median.

### Mathematical modeling of TME dynamics

We updated our previously published stochastic delay differential equation model^41^ to accurately reflect an established tumor undergoing treatment by adjusting relevant parameters and initial conditions. We now consider a stochastic differential equation model with the same four cell populations (tumor cells, MDSCs, NK cells, CTLs) with the following changes:

- We set τ = 0 (as the MDSC delay to the microenvironment is no longer relevant),
- We initialize cell populations at the numerically determined steady state of the established tumor rather than starting with only a couple of tumor cells,
- We implement parameter changes to match population sizes from the single-cell murine data,
- We implement additive noise instead of multiplicative noise.

Therefore, our modified model has the same mathematical form^41^, but with parameter values more accurately reflecting the microenvironment under consideration. Parameter changes are also shown in **Figure 6**. A full description of parameters is shown in **Supplementary Table 3**.

We incorporate treatment in the model by the modulation of important parameters that represent interactions between cell populations within the tumor microenvironment (**Figure 6A**). The relative scaling of parameters is determined by CellChat interaction scores^31^. Testing many different treatment effects and the influence of predicted changes in signaling is simple in the context of the model, but is not feasible in an experimental laboratory setting, as the combination of parameters/LR pairs/treatments/etc is combinatorially huge. We first identified five parameters in the mathematical model that are likely to be influenced by treatment, namely inhibition and stimulation rates. These parameters are β_1_ (tumor inhibition by NK cells), β_2_ (tumor inhibition by CTL cells), β_3_ (NK inhibition by MDSCs), β_4_ (CTL inhibition by MDSCs), and α_6_ (CTL stimulation by tumor-NK interaction). Through extensive CellChat/CellCircuit analysis of relevant LR pairs and pathways between these cell populations, we determined two main treatment changes under EPC treatment, namely that MDSC inhibition of CTL cells is predicted to decrease (β_4_ → β_4_ × 0.38) and CTL stimulation is predicted to increase (α_6_ → α_6_ × 1.4). For patient R-02, we observe a dramatic increase in plasma cells, which can lead to increased NK cell activation to target and destroy tumor cells (β_1_ → β_1_ × 2.0).

### scRNA-seq (primary tumors)

The scRNA-seq dataset for primary tumors was obtained from Sidiropoulos *et al*.^12^ For library preparation, 10× Genomics Chromium Single-Cell 3′ RNA-seq kits v2 were used. Gene expression libraries were prepared according to the manufacturer’s protocol. RNA was extracted from 20 whole tumors from the following groups: Vehicle control (V), entinostat-treated (E), entinostat and anti-PD-1 (EP), entinostat and anti-CTLA-4 (EC), entinostat with anti-PD-1 and anti-CTLA-4 (EPC), with four biological replicates from each of the five experimental groups. Tumors were sequenced in four batches: RunA (eight tumors; 1 E, 2 EP, 2 EC, 2 EPC, 1 V), RunB (eight tumors; 1 E, 2 EP, 2 EC, 2 EPC, 1 V), Pilot1 (two tumors, 1 E and 1 V), and Pilot2 (two tumors, 1 E and 1 V). Each batch had an approximately equal assortment of samples from each treatment group to reduce technical biases. Illumina HiSeqX Ten or NovaSeq was used to generate approximately 6.5 billion total reads. Paired-end reads were processed using CellRanger v3.0.2 and mapped to the mm10 transcriptome v1.2.0 by 10× Genomics with default settings. ScanPy v1.9.1 and Python v3 were used for quality control and basic filtering. For gene filtering, all genes expressed in fewer than 3 cells within a tumor were removed. Cells expressing fewer than 200 genes or more than 8,000 genes or having more than 15% mitochondrial gene expression were also removed. Gene expression was total-count normalized to 10,000 reads per cell and log transformed. Highly variable genes were identified using default ScanPy parameters, and the total counts per cell and the percentage of mitochondrial genes expressed were regressed out. Finally, gene expression was scaled to unit variance, and values exceeding 10 standard deviations were removed. There were 54,636 cells and 19,606 genes after pre- processing. Batch effects were corrected using the ComBat batch correction package. Neighborhood graphs were constructed using 10 nearest neighbors and 30 principal components. Cells were clustered together using Louvain clustering, and 6 main clusters were identified (with resolution parameter 0.1).

### scRNA-seq (lung mets)

For library preparation, 10× Genomics Chromium Single-Cell 3′ RNA-seq kits v3 were used. Gene expression libraries were prepared according to the manufacturer’s protocol. Each batch had an approximately equal assortment of samples from each treatment group to reduce technical biases. Illumina HiSeqX Ten or NovaSeq was used to generate approximately 6.5 billion total reads. Paired-end reads were processed using CellRanger v4.0.0 and mapped to the mm10 transcriptome v3.0.0 by 10× Genomics with default settings. ScanPy v1.9.1 and Python v3 were used for quality control and basic filtering. For gene filtering, all genes expressed in fewer than 3 cells within a sample were removed. Cells expressing fewer than 200 genes or more than 8,000 genes or having more than 15% mitochondrial gene expression were also removed. Gene expression was normalized by total count to 10,000 reads per cell and log-transformed. Highly variable genes were identified using default ScanPy parameters, and the total counts per cell and the percentage of mitochondrial genes expressed were regressed out. Finally, gene expression was scaled to unit variance, and values exceeding 10 standard deviations were removed. There were 58,097 cells and 21,007 genes after pre-processing. Batch effects were corrected using the ComBat batch correction package. Neighborhood graphs were constructed using 10 nearest neighbors and 30 principal components. Cells were clustered together using Louvain clustering, and 8 main clusters were identified (with resolution parameter 0.1).

The following main clusters: cancer, mature myeloid, MDSCs, T cells, B cells, and NK cells were subclustered similarly as above (see **Figures 2, S2-S6**). Subclusters that likely represented doublets were removed from the analysis.

### Cell proportion test

We performed cell proportion analysis using the R package propeller t-tests with logit transformation^83^ and simple t-tests to compare treatment V with other treatment groups. The analysis focused exclusively on immune cell populations, and the proportions used in the tests were calculated relative to the total immune cell population within each treatment group. False Discovery Rate (FDR) correction was applied to account for multiple testing.

### CellChat analysis

We implement the CellChat R package on the lung mets scRNA seq data with different treatments (V, E, EPC, PC) independently. We have three different modules: CC-model, CC-suppression and CC-DC, which include different cell populations as input to CellChat (**Figure 4A**). CellChat results are included in the Supplementary Information in tabular format.

### CellCircuit analysis

To emphasize the key interactions, weaker interactions were filtered out. In the CC-suppression, cell state interactions were ranked by strength, retaining the top 50% for downstream analysis, resulting in the removal of approximately 10% of weighted edges in each interaction network. For the CC-model module, a fixed absolute cutoff of 0.293 was applied across all treatments, corresponding to the top two-thirds of interactions in the V-treated cells. We saw an increase in the number of highly weighted interactions in this comparison with E or EPC, which would lead to throwing away many more interactions at a percentile cutoff, so we use the absolute value cutoff that is set by V.

From the filtered interaction matrix for each module with V, a weighted directed graph was constructed. Within the weighted directed graph, 2-cell and 3-cell motifs were identified by IGLADFindSubisomorphisms function from Mathematica package IGraphM^83^. The average weight per edge was summed up within each subgraph allowing comparison of different subgraph classes.

To assess the statistical significance of the identified motif patterns in V, 10,000 random networks were generated by rewiring the connections within the original weighted directed graph using the IGRewire function. This rewiring process preserves the in-degree and out-degree of each node while randomly permuting the interaction weights, allowing self-edges. In each randomized network, the same subgraph scoring procedure was applied as in the original network. Using the score distribution from these randomized networks, Z-scores were calculated for the subgraphs identified in the original network, with a significance cutoff set at 1.6^42^.

For each module, we independently selected the most enriched motifs from two- and three-state configurations (**Figure 4C**). In the CC-suppression module, fully connected two- and three-state motifs were chosen (**Figure 4D; S9C**). In the CC-model module, unidirectional two-state motifs and feedforward/feedback three-state motifs were selected (**Figure 4E; S10B**). After identifying significantly enriched motifs in each module under treatment V, motif instances within each class were ranked by total interaction weight. To further refine our analysis, we concentrated on circuits involving interactions between distinct cell states, excluding intra-state signaling such as macrophage-to-neutrophil interactions. This refinement resulted in 10 two-cell and 10 three-cell instances for CC-suppression, and 5 two-cell and 4 three-cell instances for CC-model. From each motif pattern, the highest-weighted instance involving biologically relevant cell-cell signaling was selected for further analysis (**Figure 4C-D**).

For each selected instance, corresponding subgraphs from other treatments (E, EPC, and PC) were generated by Mathematica to enable treatment-wise comparisons (**Figure 4F; S10E**). In the CellChat output, each edge represents signaling strength across pathways and ligand-receptor pairs. Within each instance, changes in interaction weight from treatment V to other treatments were quantified by taking the logarithm of the fraction of strength in treatment and strength in V, providing a detailed view of pathway- and ligand-receptor-specific differences. We excluded the pathways with a strength smaller than 0.01 in all treatments. (**S9 D-F; S10 D**).

### Analysis of treatment effect on tumor and tumor heterogeneity

There are 3098 tumor cells after 574 doublets were removed. Batch effects were corrected by the ComBat batch correction package incorporated in ScanPy v1.9.1. Highly variable genes are selected by default parameters in ScanPy. 30 PCs and 30 nearest neighbors are used to create a neighborhood graph. Tumor cell sub-states were clustered together using Louvain clustering with a resolution of 0.175, and 3 main clusters were identified. Cell cycle genes^84^ were scored to infer cell cycle phase by function in ScanPy. We used propeller^85^ with default parameters to evaluate whether treatment induced significant proportional changes across both subclusters and cell cycle phases, with a significant threshold FDR smaller than 0.05.

To quantitatively evaluate the treatment effect, linear discriminant analysis (LDA) was performed pair-wisely in R with genes significantly differentially expressed (alpha = 0.5) between vehicle and treatments E, EPC, and PC by Wilcoxon rank sum test in ScanPy. 10 genes with the highest LDA coefficients were selected and used to create LDA embeddings for tumor cells under vehicle versus treatments by using 80% of the cells as a training dataset. The degree of shifts between untreated and treated cells in LDA embeddings was quantified via the Earth mover’s distance (EMD)^86^ via the emdist R package^87^. Classification accuracy by LDA on the test dataset was also obtained. Logistic regression was performed, and the top ten genes with the highest coefficient were selected. Moreover, logistic regression with regularization was performed iteratively on gene expression in Python with the scikit-learn package, comparing E, EPC, and PC with V, as described by Liu *et al*^88^. In each iteration, 20% of genes are removed until the top 10 genes are left. Ten runs of such iterative logistic regression were performed, and genes appearing more than five times were selected. Hallmark Epithelial Mesenchymal transition markers^89^, stem marker genes^90–92^, and PAM50 with subtypes^93^ were scored independently on treatments by using the UCell R package^94^. Pairwise t-tests were performed on each score to evaluate treatment effects.

**Table 1.**
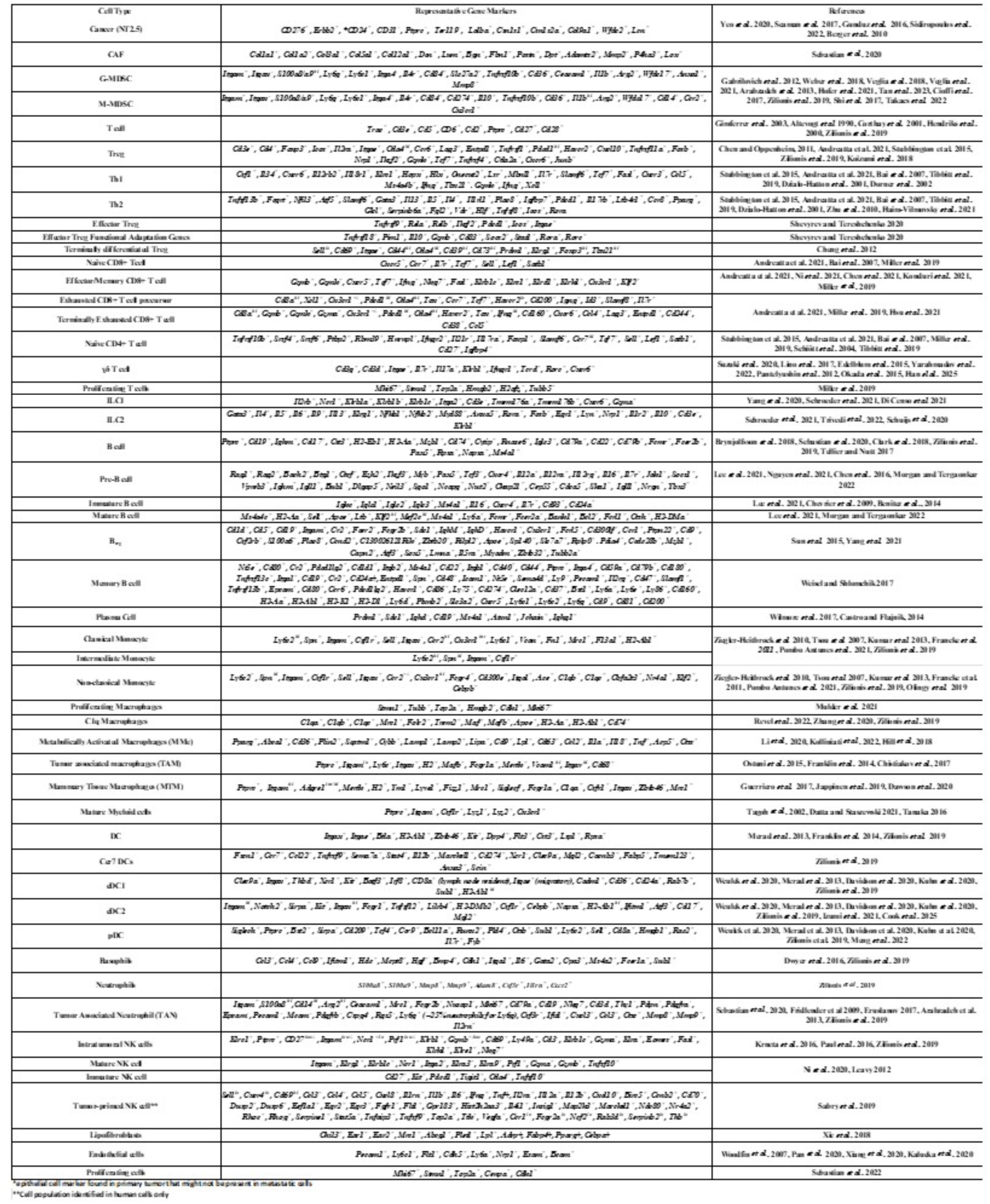
Curated gene list used for scRNA-seq cluster annotations. Signature marker genes were collected from various sources outlined in the “References” column, prioritizing articles describing single-cell RNA sequencing experiments using mouse models of cancer. Complete references found in Supplementary File 1.

**Table 2.**
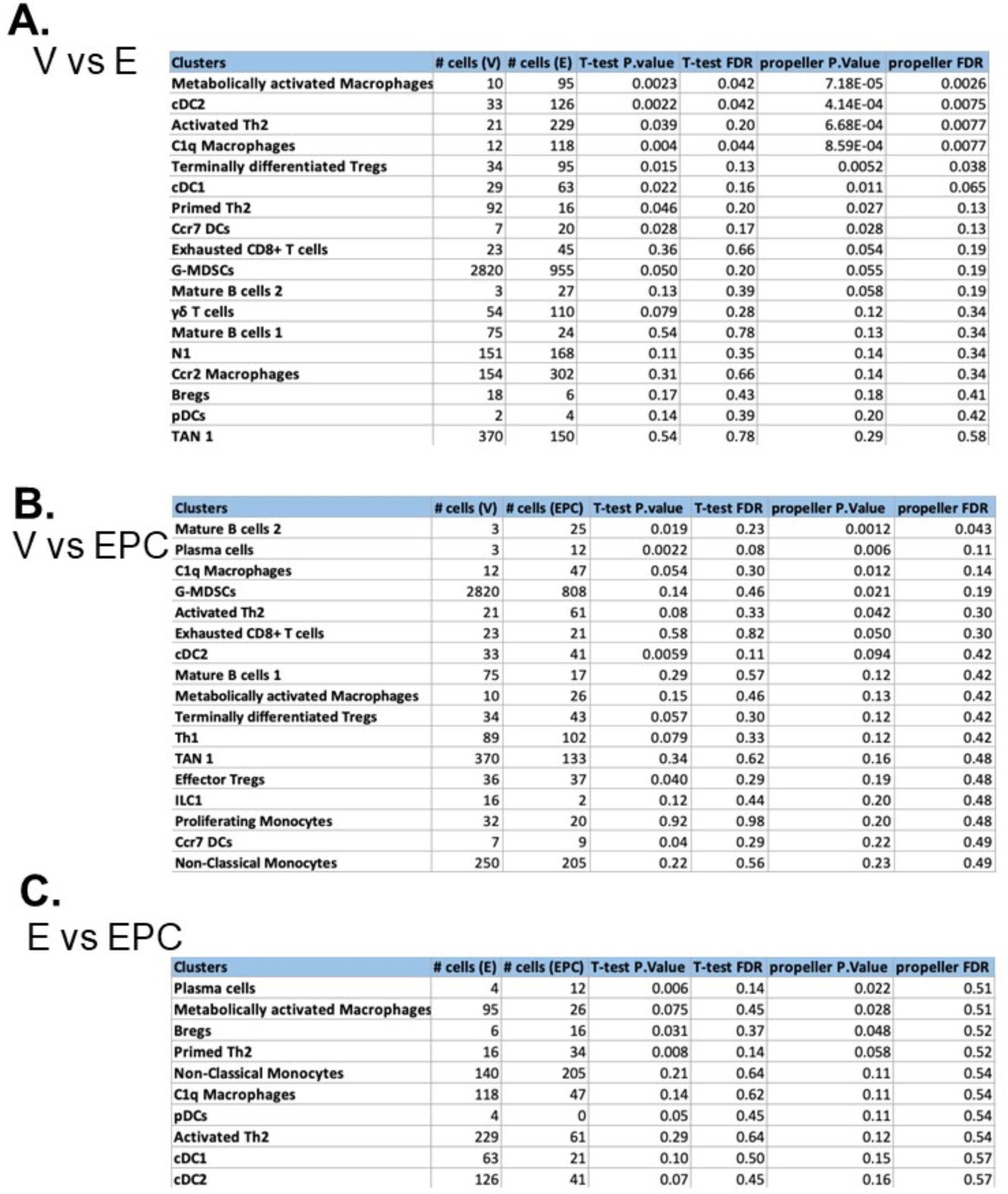
Immune cell population proportional test comparing treatments pair-wisely (A: V vs E; B: V vs EPC; C: E vs EPC). Each comparison was performed by t-test and R package propeller t-test with logit transformation. All p-values were corrected by FDR and immune cell populations with propeller FDR < 0.6 were shown in each table.

A. V vs E
| Clusters | # cells (V) | # cells (E) | T-test P.value | T-test FDR | propeller P.Value | propeller FDR |
| --- | --- | --- | --- | --- | --- | --- |
| Metabolically activated Macrophages | 10 | 95 | 0.0023 | 0.042 | 7.18E-05 | 0.0026 |
| cDC2 | 33 | 126 | 0.0022 | 0.042 | 4.14E-04 | 0.0075 |
| Activated Th2 | 21 | 229 | 0.039 | 0.20 | 6.68E-04 | 0.0077 |
| C1q Macrophages | 12 | 118 | 0.004 | 0.044 | 8.59E-04 | 0.0077 |
| Terminally differentiated Tregs | 34 | 95 | 0.015 | 0.13 | 0.0052 | 0.038 |
| cDC1 | 29 | 63 | 0.022 | 0.16 | 0.011 | 0.065 |
| Primed Th2 | 92 | 16 | 0.046 | 0.20 | 0.027 | 0.13 |
| Ccr7 DCs | 7 | 20 | 0.028 | 0.17 | 0.028 | 0.13 |
| Exhausted CD8+ T cells | 23 | 45 | 0.36 | 0.66 | 0.054 | 0.19 |
| G-MDSCs | 2820 | 955 | 0.050 | 0.20 | 0.055 | 0.19 |
| Mature B cells 2 | 3 | 27 | 0.13 | 0.39 | 0.058 | 0.19 |
| γδ T cells | 54 | 110 | 0.079 | 0.28 | 0.12 | 0.34 |
| Mature B cells 1 | 75 | 24 | 0.54 | 0.78 | 0.13 | 0.34 |
| N1 | 151 | 168 | 0.11 | 0.35 | 0.14 | 0.34 |
| Ccr2 Macrophages | 154 | 302 | 0.31 | 0.66 | 0.14 | 0.34 |
| Bregs | 18 | 6 | 0.17 | 0.43 | 0.18 | 0.41 |
| pDCs | 2 | 4 | 0.14 | 0.39 | 0.20 | 0.42 |
| TAN 1 | 370 | 150 | 0.54 | 0.78 | 0.29 | 0.58 |

B. V vs EPC
| Clusters | # cells (V) | # cells (EPC) | T-test P.value | T-test FDR | propeller P.Value | propeller FDR |
| --- | --- | --- | --- | --- | --- | --- |
| Mature B cells 2 | 3 | 25 | 0.019 | 0.23 | 0.0012 | 0.043 |
| Plasma cells | 3 | 12 | 0.0022 | 0.08 | 0.006 | 0.11 |
| C1q Macrophages | 12 | 47 | 0.054 | 0.30 | 0.012 | 0.14 |
| G-MDSCs | 2820 | 808 | 0.14 | 0.46 | 0.021 | 0.19 |
| Activated Th2 | 21 | 61 | 0.08 | 0.33 | 0.042 | 0.30 |
| Exhausted CD8+ T cells | 23 | 21 | 0.58 | 0.82 | 0.050 | 0.30 |
| cDC2 | 33 | 41 | 0.0059 | 0.11 | 0.094 | 0.42 |
| Mature B cells 1 | 75 | 17 | 0.29 | 0.57 | 0.12 | 0.42 |
| Metabolically activated Macrophages | 10 | 26 | 0.15 | 0.46 | 0.13 | 0.42 |
| Terminally differentiated Tregs | 34 | 43 | 0.057 | 0.30 | 0.12 | 0.42 |
| Th1 | 89 | 102 | 0.079 | 0.33 | 0.12 | 0.42 |
| TAN 1 | 370 | 133 | 0.34 | 0.62 | 0.16 | 0.48 |
| Effector Tregs | 36 | 37 | 0.040 | 0.29 | 0.19 | 0.48 |
| ILC1 | 16 | 2 | 0.12 | 0.44 | 0.20 | 0.48 |
| Proliferating Monocytes | 32 | 20 | 0.92 | 0.98 | 0.20 | 0.48 |
| Ccr7 DCs | 7 | 9 | 0.04 | 0.29 | 0.22 | 0.49 |
| Non-Classical Monocytes | 250 | 205 | 0.22 | 0.56 | 0.23 | 0.49 |

| Clusters | # cells (E) | # cells (EPC) | T-test P.Value | T-test FDR | propeller P.Value | propeller FDR |
| --- | --- | --- | --- | --- | --- | --- |
| Plasma cells | 4 | 12 | 0.006 | 0.14 | 0.022 | 0.51 |
| Metabolically activated Macrophages | 95 | 26 | 0.075 | 0.45 | 0.028 | 0.51 |
| Bregs | 6 | 16 | 0.031 | 0.37 | 0.048 | 0.52 |
| Primed Th2 | 16 | 34 | 0.008 | 0.14 | 0.058 | 0.52 |
| Non-Classical Monocytes | 140 | 205 | 0.21 | 0.64 | 0.11 | 0.54 |
| C1q Macrophages | 118 | 47 | 0.14 | 0.62 | 0.11 | 0.54 |
| pDCs | 4 | 0 | 0.05 | 0.45 | 0.11 | 0.54 |
| Activated Th2 | 229 | 61 | 0.29 | 0.64 | 0.12 | 0.54 |
| cDC1 | 63 | 21 | 0.10 | 0.50 | 0.15 | 0.57 |
| cDC2 | 126 | 41 | 0.07 | 0.45 | 0.16 | 0.57 |

**Table 3.** Table showing mathematical model parameters and how they change with treatment.

| Notation | Description | CIR | Data | Change | Treatment |
| --- | --- | --- | --- | --- | --- |
| $x_T(t), t \leq 0$ | initial condition for tumor cells | 1 or 2 | steady state | yes | - |
| $x_{\text{MDSC}}(0)$ | initial condition for MDSCs | $\alpha_2/\zeta_2$ | steady state | yes | - |
| $x_{\text{NK}}(0)$ | initial condition for NK cells | $\frac{\zeta_2 \alpha_4}{\alpha_2 \beta_3 + \zeta_2 \zeta_3}$ | steady state | yes | - |
| $x_{\text{CTL}}(0)$ | initial condition for CTL cells | 0 | steady state | yes | - |
| $\tau$ | delay parameter for MDSCs | varies | 0 | - | - |
| $\alpha_1$ | tumor growth rate | $10^{-1}$ | $10^{-1}$ | - | - |
| $\eta$ | tumor maximum size | $10^7$ | $10^7$ | - | - |
| $\beta_1$ | tumor cells inhibition rate by NK cells | $3.5 \times 10^{-6}$ | $6 \times 10^{-4}$ | yes | no change |
| $\beta_2$ | tumor cells inhibition rate by CTL cells | $1.1 \times 10^{-7}$ | $7 \times 10^{-4}$ | yes | no change |
| $\zeta_1$ | tumor cell death rate | 0 | 0 | - | - |
| $\alpha_2$ | MDSCs circulating rate | $10^2$ | $5 \times 10^2$ | yes | - |
| $\alpha_3$ | MDSCs expansion coefficient | $10^8$ | $10^8$ | - | - |
| $\zeta_2$ | MDSCs death rate | 0.2 | 0.2 | - | - |
| $\alpha_4$ | NK cells circulating rate | $1.4 \times 10^4$ | $5 \times 10^1$ | yes | - |
| $\alpha_5$ | NK cells expansion coefficient | $2.5 \times 10^{-2}$ | $2.5 \times 10^{-2}$ | - | - |
| $\beta_3$ | NK cells inhibition rate by MDSCs | $4 \times 10^{-5}$ | $4 \times 10^{-5}$ | - | no change |
| $\zeta_3$ | NK cells death rate | $4.12 \times 10^{-2}$ | $4.12 \times 10^{-2}$ | - | - |
| $\alpha_6$ | CTL stimulation by tumor-NK cell interaction | $10^3$ | $4 \times 10^{-3}$ | yes | $\alpha_6 \rightarrow \alpha_6 \times 1.4$ |
| $\alpha_7$ | CTL expansion coefficient | $10^{-1}$ | $10^{-1}$ | - | - |
| $\beta_4$ | CTL inhibition rate by MDSCs | $10^{-4}$ | $10^{-4}$ | - | $\beta_4 \rightarrow \beta_4 \times 0.38$ |
| $\zeta_4$ | CTL death rate | $2 \times 10^{-2}$ | $2 \times 10^{-2}$ | - | - |
| $\gamma_1$ | steepness of MDSC production | $10^{10}$ | $10^{10}$ | - | - |
| $\gamma_2$ | steepness of NK production | $2.02 \times 10^7$ | $2.02 \times 10^7$ | - | - |
| $\gamma_3$ | steepness of CTL production | $2.02 \times 10^7$ | $2.02 \times 10^7$ | - | - |

## Author Contributions

Conceptualization: EG, JK, YL, ALM, ERT.

Methodology: EG, JK, YL, AB, AGB, JE, VS, RMC, WH, ALM, ERT.

Software: EG, JK, YL, XW, ALM.

Investigation: EG, JK, YL, XW, AB, AGB, BAZ, JJ, SMS, WH.

Formal analysis: JK, YL, XW, ALM.

Resources: WH, JE, VS, RMC

Writing: original draft: EG, JK, ALM, ERT.

Supervision: ALM, ERT.

All authors read and approved the final manuscript:

## Acknowledgements

We would like to acknowledge the patients and their families who participated in NCI-9844 without whom the translational relevance would not be possible. We would also like to thank the site PIs from NCI-9844: Adam Brufsky, Patricia LoRusso, Vincent Chung, and Yuan Yuan. We would also like to acknowledge others who were part of the original team for NCI-9844 who contributed to ensuring completion of the trial and provided advice and storage of specimens and personnel support for access to clinical outcomes data including Richard Piekarz, Howard Streicher, Elizabeth M. Jaffee, and Ashley Cimino-Mathews. As well as James Leatherman, Christine Rafie, Melinda Downs, and Ashley O’Connor, who contributed to biospecimen and data management. We also would like to thank the core facilities at USC Norris Comprehensive Cancer Center including the Molecular Genomics Core, Translational Pathology Core, Flow Cytometry and Immune Monitoring Core, and Stem Cell Flow Cytometry Core and was supported by award number P30CA014089 from the National Cancer Institute. The content is solely the responsibility of the authors and does not necessarily represent the official views of the National Cancer Institute or the National Institutes of Health This study was also supported by the NCI R01CA283169 (ERT), Tower Cancer Research Foundation (ERT), American Association for Cancer Research and the Breast Cancer Research Foundation Grant (ERT), METAVivor Foundation (ERT), Wright Foundation (ERT), Concern Foundation (ERT), NIGMS R35GM143019 (ALM), NSF DMS 2045327 (ALM) and the V Foundation (Translational Award 2017 to RMC and VS). JE is supported by a K08 grant from the NCI (K08CA245220).

