## Supplementary Figures for "Cancer systems immunology reveals myeloid—T cell interactions and B cell activation mediate response to checkpoint inhibition in metastatic breast cancer"

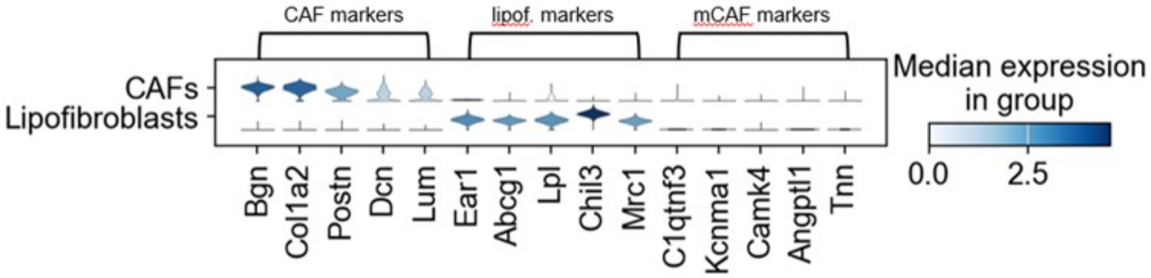
CAF markers
lipof. markers
mCAF markers
CAFs
Lipofibroblasts
Median expression
in group
0.0
2.5
Bgn
Col1a2
Postn
Dcn
Lum
Ear1
Abcg1
Lpl
Chil3
Mrc1
C1qtnf3
Kcnma1
Camk4
Angptl1
Tnn

**S1**

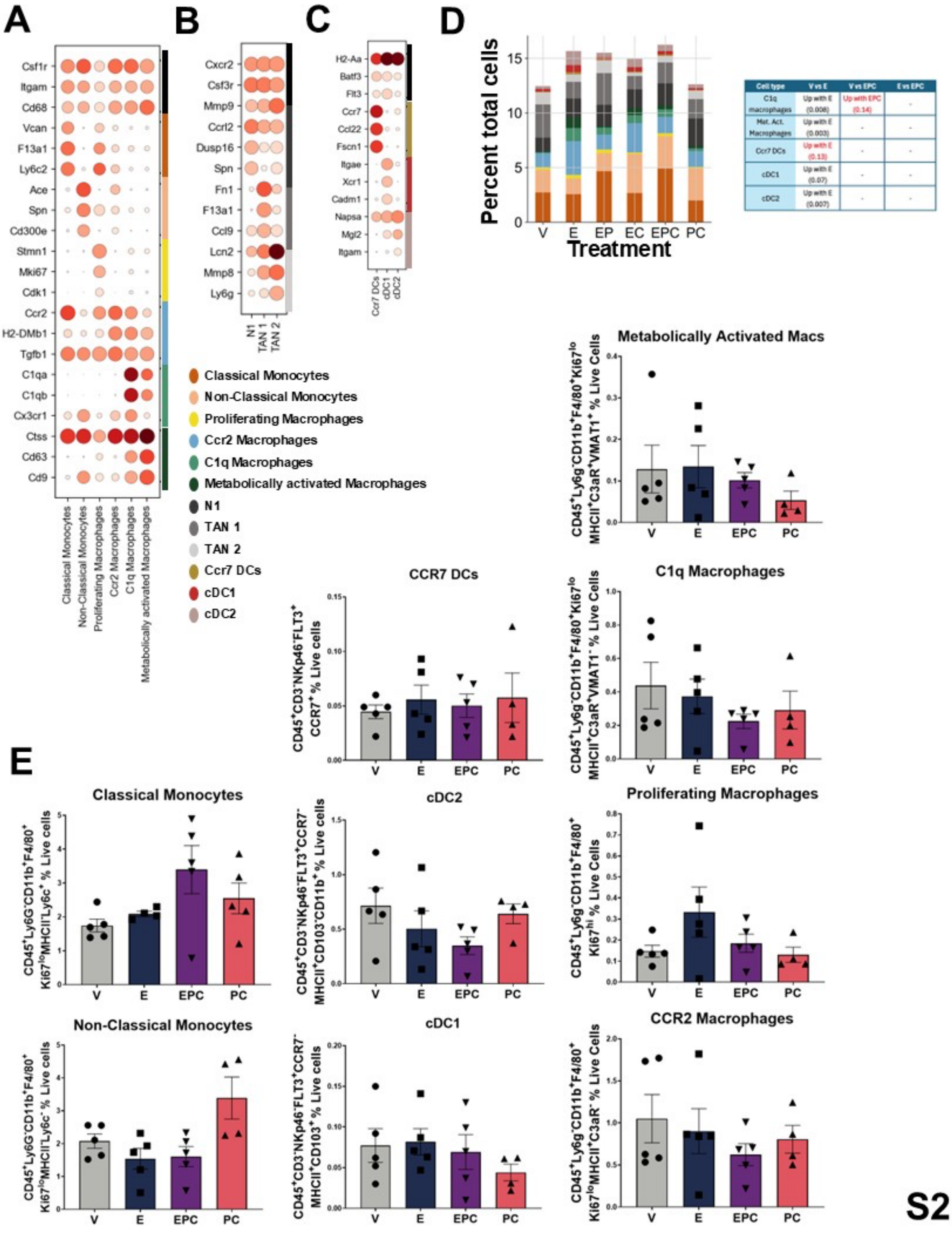


| Cell type | V vs E | V vs EPC | E vs EPC |
| --- | --- | --- | --- |
| C1q macrophages | Up with E (0.008) | Up with EPC (0.14) | - |
| Met. Act. Macrophages | Up with E (0.003) | - | - |
| Ccr7 DCs | Up with E (0.13) | - | - |
| cDC1 | Up with E (0.07) | - | - |
| cDC2 | Up with E (0.007) | - | - |

S2

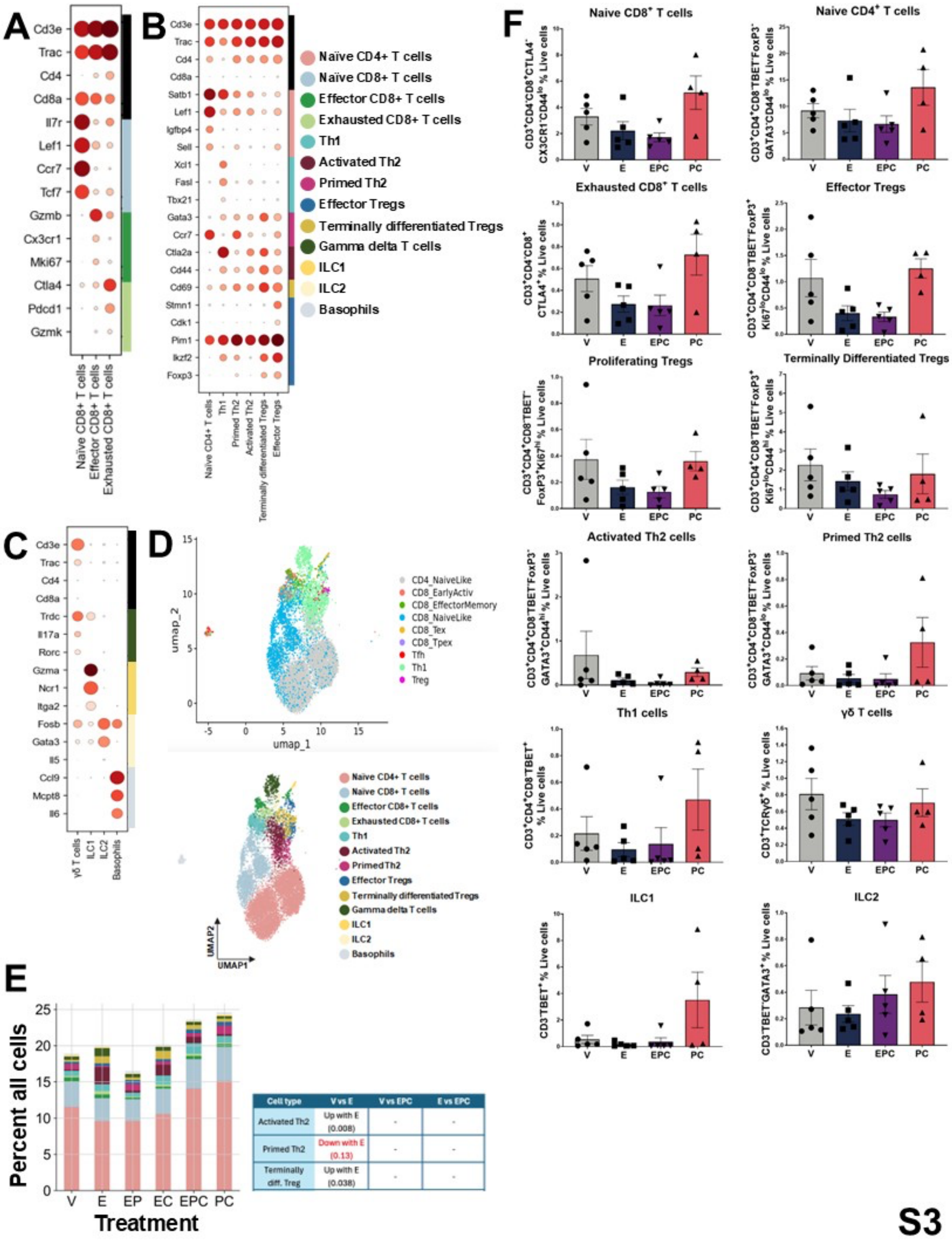


| Cell type | V vs E | V vs EPC | E vs EPC |
|---|---|---|---|
| Activated Th2 | Up with E (0.008) | - | - |
| Primed Th2 | Down with E (0.13) | - | - |
| Terminally diff. Treg | Up with E (0.038) | - | - |

**S3**

## 4T1

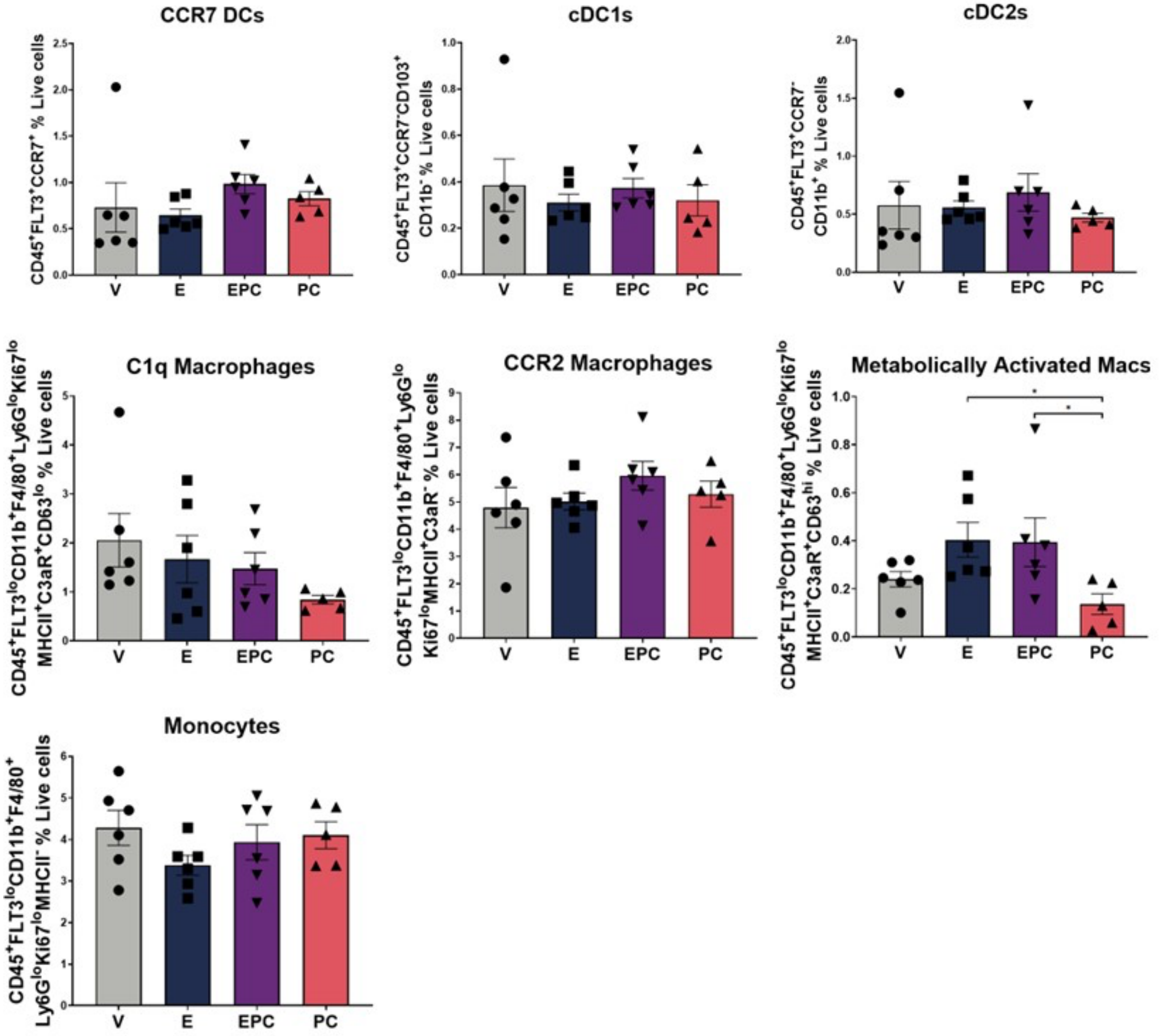
CCR7 DCs
CD45+FLT3+CCR7+ % Live cells
cDC1s
CD45+FLT3+CCR7-CD103+ CD11b- % Live cells
cDC2s
CD45+FLT3+CCR7- CD11b+ % Live cells
C1q Macrophages
CD45+FLT3loCD11b+F4/80+Ly6GloKi67lo MHCII+C3aR+CD63lo % Live cells
CCR2 Macrophages
CD45+FLT3loCD11b+F4/80+Ly6Glo Ki67loMHCII+C3aR- % Live cells
Metabolically Activated Macs
CD45+FLT3loCD11b+F4/80+Ly6GloKi67lo MHCII+C3aR+CD63hi % Live cells
Monocytes
CD45+FLT3loCD11b+F4/80+ Ly6GloKi67loMHCII- % Live cells
V
E
EPC
PC

**S4**

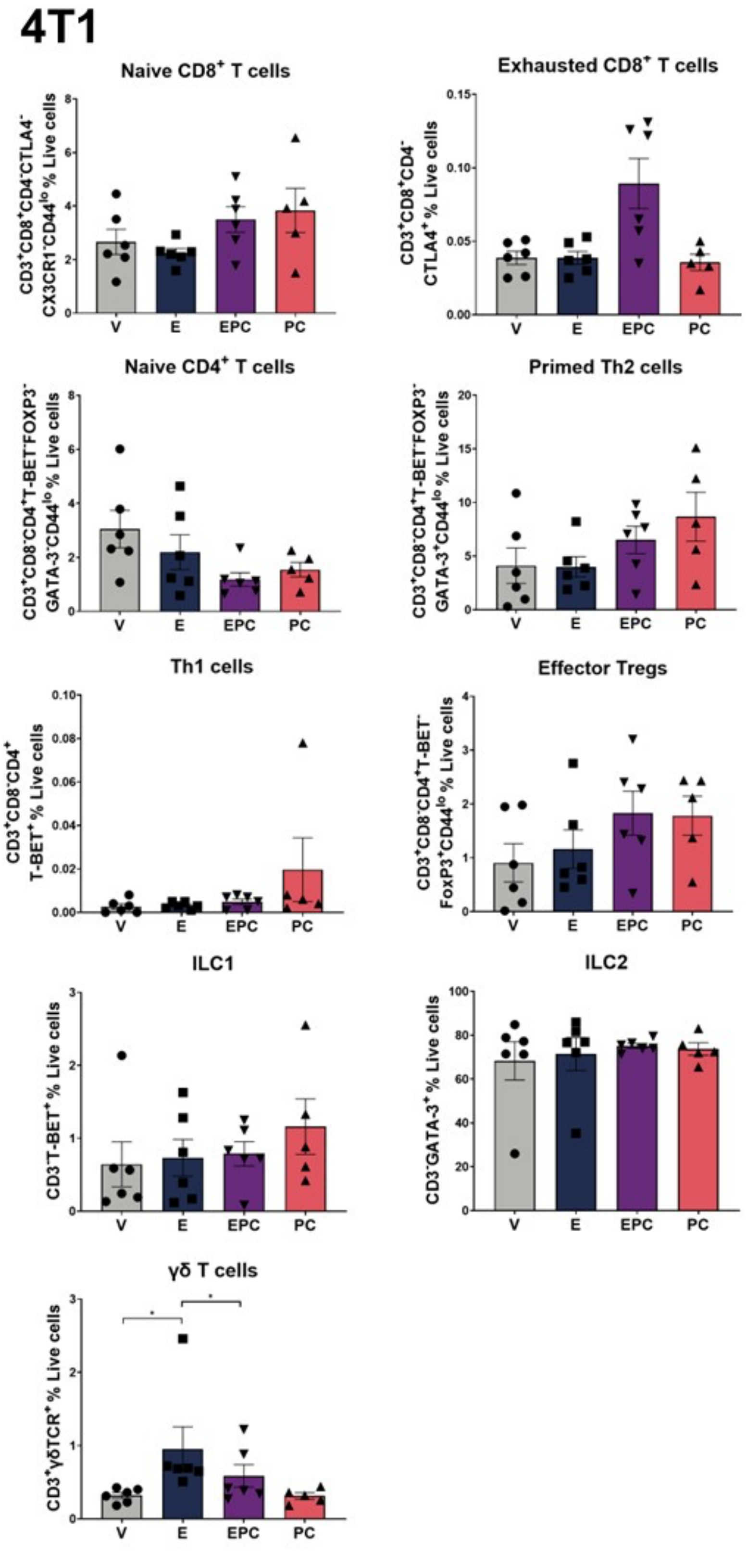
4T1
Naive CD8+ T cells
CD3+CD8+CD4-CTLA4- CX3CR1-CD44lo % Live cells
Exhausted CD8+ T cells
CD3+CD8+CD4- CTLA4+ % Live cells
Naive CD4+ T cells
Primed Th2 cells
Th1 cells
CD3+CD8-CD4+ T-BET+ % Live cells
Effector Tregs
ILC1
CD3-T-BET+ % Live cells
ILC2
CD3-GATA-3+ % Live cells
γδ T cells
CD3+γδTCR+ % Live cells
V
E
EPC
PC

S5

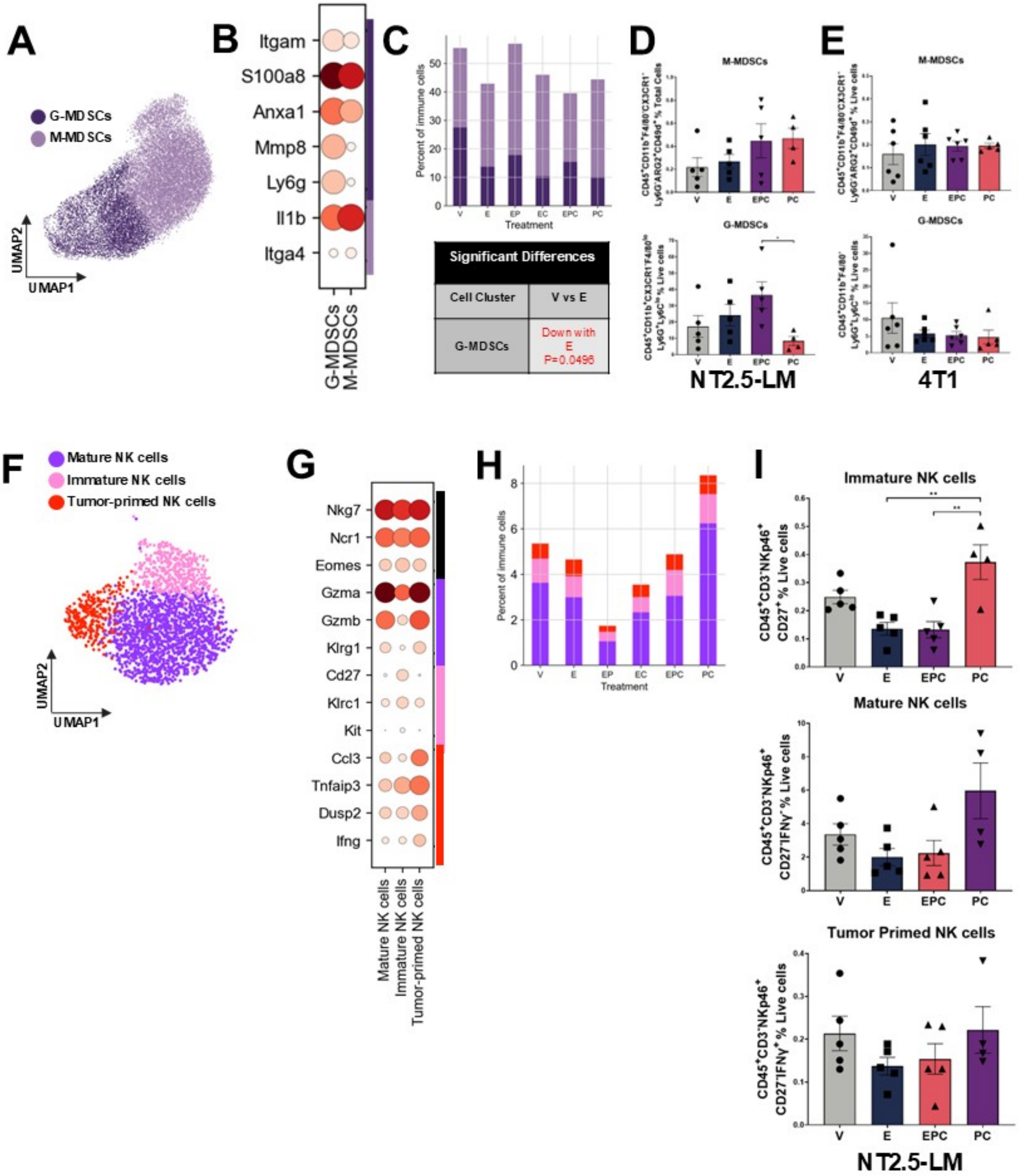
A
G-MDSCs
M-MDSCs
UMAP2
UMAP1
B
Itgam
S100a8
Anxa1
Mmp8
Ly6g
Il1b
Itga4
G-MDSCs
M-MDSCs
C
Percent of immune cells
V
E
EP
EC
EPC
PC
Treatment
Significant Differences
Cell Cluster
V vs E
G-MDSCs
Down with E P=0.0496
D
M-MDSCs
G-MDSCs
NT2.5-LM
E
M-MDSCs
G-MDSCs
4T1
F
Mature NK cells
Immature NK cells
Tumor-primed NK cells
UMAP2
UMAP1
G
Nkg7
Ncr1
Eomes
Gzma
Gzmb
Klrg1
Cd27
Klrc1
Kit
Ccl3
Tnfaip3
Dusp2
Ifng
Mature NK cells
Immature NK cells
Tumor-primed NK cells
H
Percent of immune cells
Treatment
I
Immature NK cells
CD45⁺CD3⁻NKp46⁺ CD27⁺ % Live cells
Mature NK cells
CD45⁺CD3⁻NKp46⁺ CD27⁻IFNγ⁻ % Live cells
Tumor Primed NK cells
CD45⁺CD3⁻NKp46⁺ CD27⁻IFNγ⁺ % Live cells
V
E
EPC
PC
NT2.5-LM

**S6**

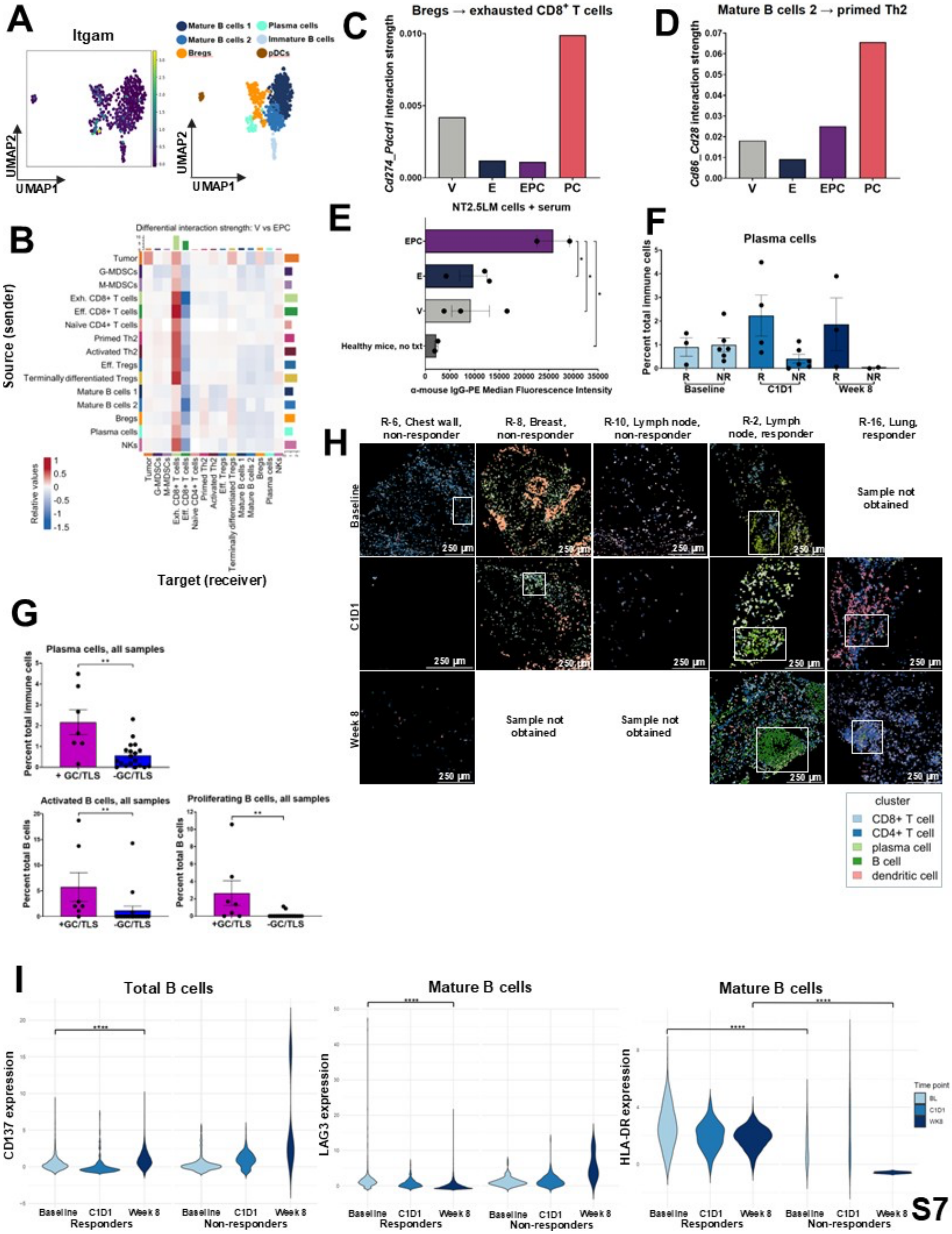
A
Itgam
UMAP2
UMAP1
Mature B cells 1
Mature B cells 2
Bregs
Plasma cells
Immature B cells
pDCs
C
Bregs → exhausted CD8+ T cells
Cd274_Pdcd1 interaction strength
V
E
EPC
PC
D
Mature B cells 2 → primed Th2
Cd86_Cd28 interaction strength
B
Differential interaction strength: V vs EPC
Source (sender)
Target (receiver)
Tumor
G-MDSCs
M-MDSCs
Exh. CD8+ T cells
Eff. CD8+ T cells
Naive CD4+ T cells
Primed Th2
Activated Th2
Eff. Tregs
Terminally differentiated Tregs
Mature B cells 1
Mature B cells 2
Bregs
Plasma cells
NKs
Relative values
E
NT2.5LM cells + serum
Healthy mice, no txt
α-mouse IgG-PE Median Fluorescence Intensity
F
Plasma cells
Percent total immune cells
Baseline
C1D1
Week 8
R
NR
G
Plasma cells, all samples
Percent total immune cells
+ GC/TLS
-GC/TLS
Activated B cells, all samples
Percent total B cells
Proliferating B cells, all samples
H
R-6, Chest wall, non-responder
R-8, Breast, non-responder
R-10, Lymph node, non-responder
R-2, Lymph node, responder
R-16, Lung, responder
Baseline
C1D1
Week 8
Sample not obtained
250 μm
cluster
CD8+ T cell
CD4+ T cell
plasma cell
B cell
dendritic cell
I
Total B cells
Mature B cells
CD137 expression
LAG3 expression
HLA-DR expression
Baseline
C1D1
Week 8
Responders
Non-responders
Time point
BL
C1D1
WK8
S7

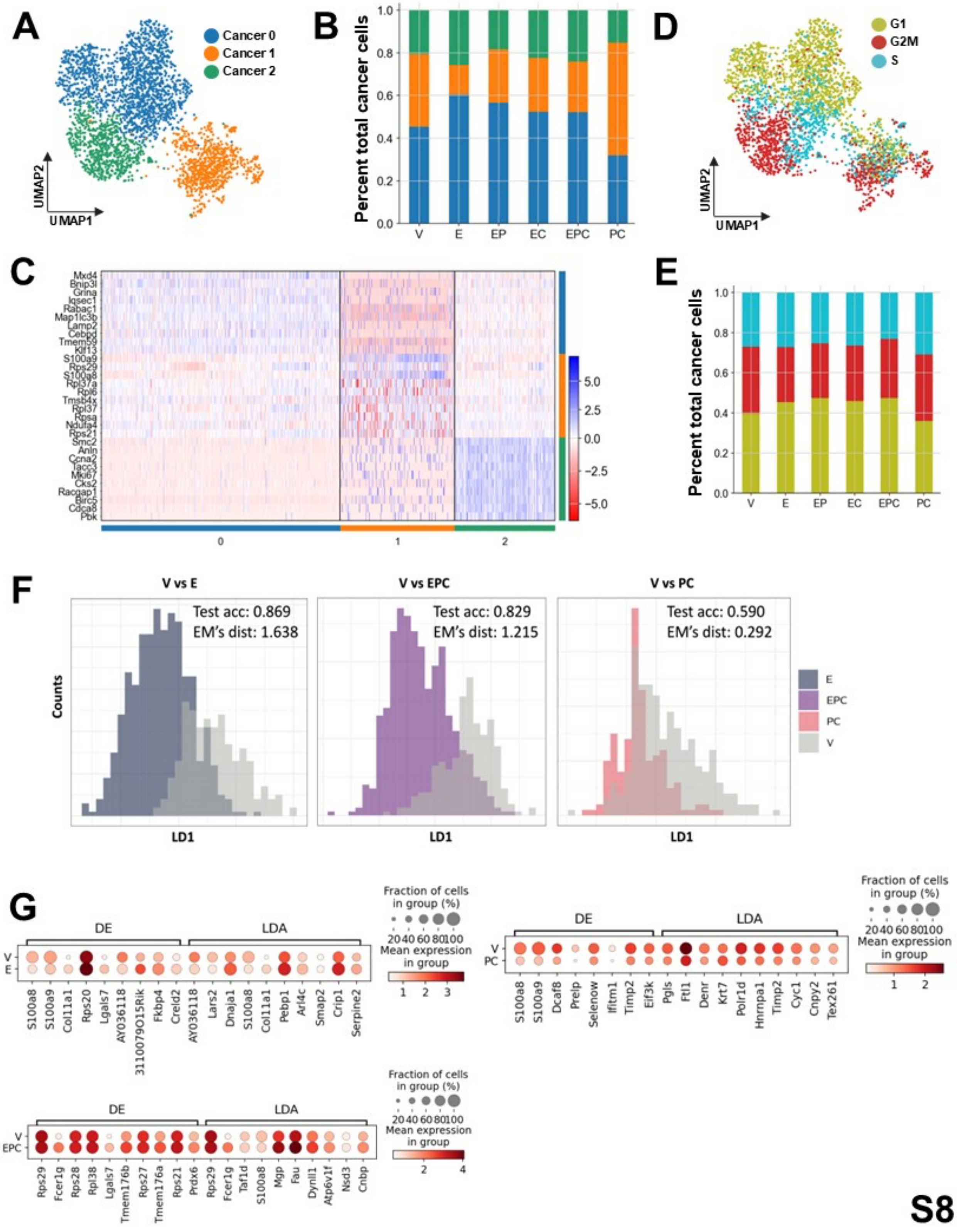
A
Cancer 0
Cancer 1
Cancer 2
UMAP2
UMAP1
B
Percent total cancer cells
V E EP EC EPC PC
D
G1
G2M
S
UMAP2
UMAP1
C
Mxd4
Bnip3l
Grina
Iqsec1
Rabac1
Map1lc3b
Lamp2
Cebpd
Tmem59
Klf13
S100a9
Rps29
S100a8
Rpl37a
Rpl6
Tmsb4x
Rpl37
Rpsa
Ndufa4
Rps21
Smc2
Anln
Ccna2
Tacc3
Mki67
Cks2
Racgap1
Birc5
Cdca8
Pbk
0 1 2
E
Percent total cancer cells
V E EP EC EPC PC
F
V vs E
Test acc: 0.869
EM's dist: 1.638
V vs EPC
Test acc: 0.829
EM's dist: 1.215
V vs PC
Test acc: 0.590
EM's dist: 0.292
Counts
LD1
E
EPC
PC
V
G
DE
LDA
Fraction of cells in group (%)
Mean expression in group
S100a8 S100a9 Col11a1 Rps20 Lgals7 AY036118 3110079O15Rik Fkbp4 Creld2 AY036118 Lars2 Dnaja1 S100a8 Col11a1 Pebp1 Arl4c Smap2 Crip1 Serpine2
S100a8 S100a9 Dcaf8 Prelp Selenow Ifitm1 Timp2 Eif3k Pgls Ftl1 Denr Krt7 Polr1d Hnrnpa1 Timp2 Cyc1 Cnpy2 Tex261
Rps29 Fcer1g Rps28 Rpl38 Lgals7 Tmem176b Rps27 Tmem176a Rps21 Prdx6 Rps29 Fcer1g Taf1d S100a8 Mgp Fau Dynll1 Atp6v1f Nsd3 Cnbp

S8

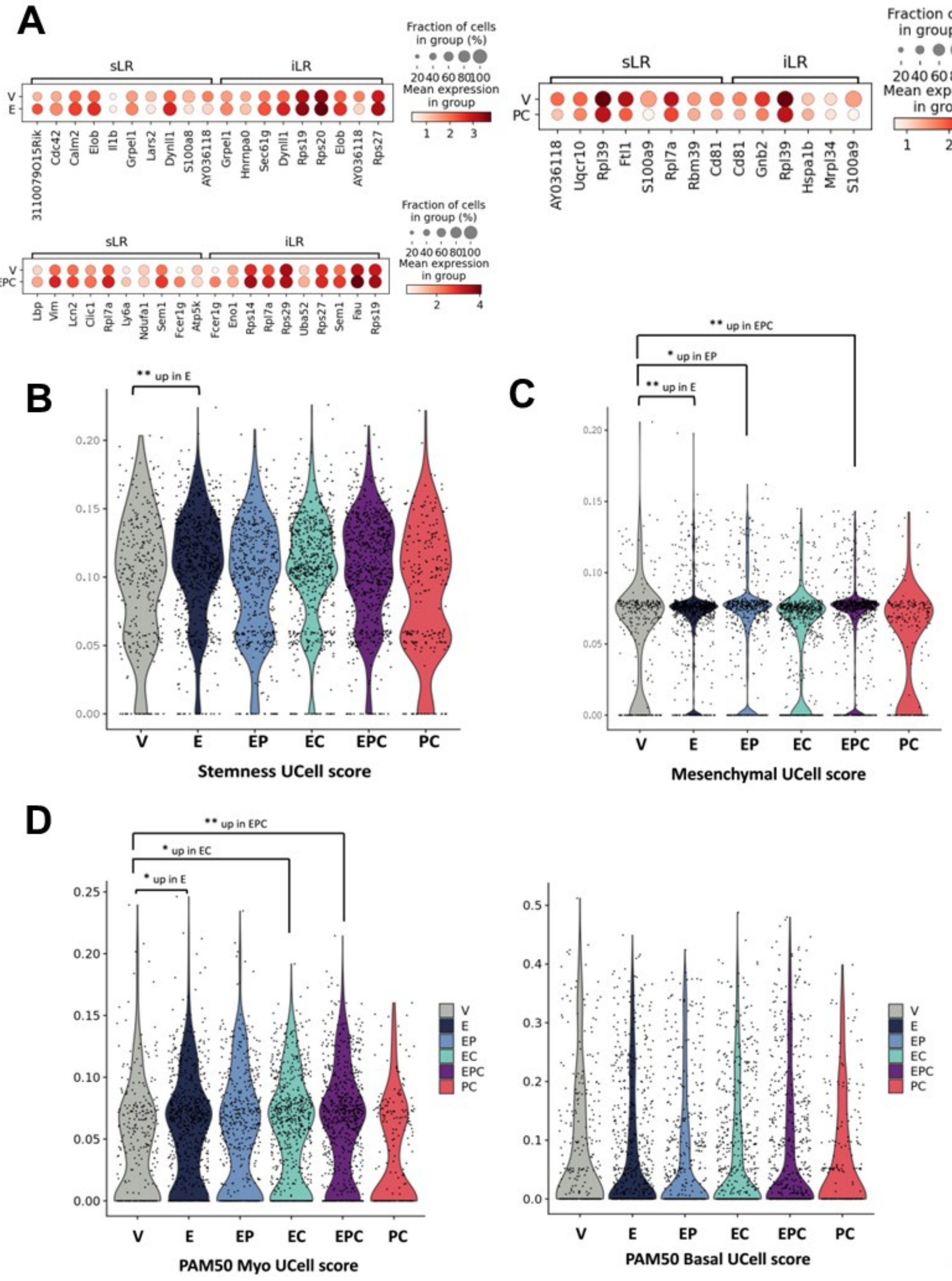
A
sLR
iLR
Fraction of cells in group (%)
20 40 60 80 100
Mean expression in group
V
E
3110079O15Rik
Cdc42
Calm2
Elob
Il1b
Grpel1
Lars2
Dynll1
S100a8
AY036118
Grpel1
Hnrnpa0
Sec61g
Dynll1
Rps19
Rps20
Elob
AY036118
Rps27
PC
AY036118
Uqcr10
Rpl39
Ftl1
S100a9
Rpl7a
Rbm39
Cd81
Cd81
Gnb2
Rpl39
Hspa1b
Mrpl34
S100a9
EPC
Lbp
Vim
Lcn2
Clic1
Rpl7a
Ly6a
Ndufa1
Sem1
Fcer1g
Atp5k
Fcer1g
Eno1
Rps14
Rpl7a
Rps29
Uba52
Rps27
Sem1
Fau
Rps19
B
** up in E
V E EP EC EPC PC
Stemness UCell score
C
** up in EPC
* up in EP
** up in E
V E EP EC EPC PC
Mesenchymal UCell score
D
** up in EPC
* up in EC
* up in E
V E EP EC EPC PC
PAM50 Myo UCell score
PAM50 Basal UCell score

S9

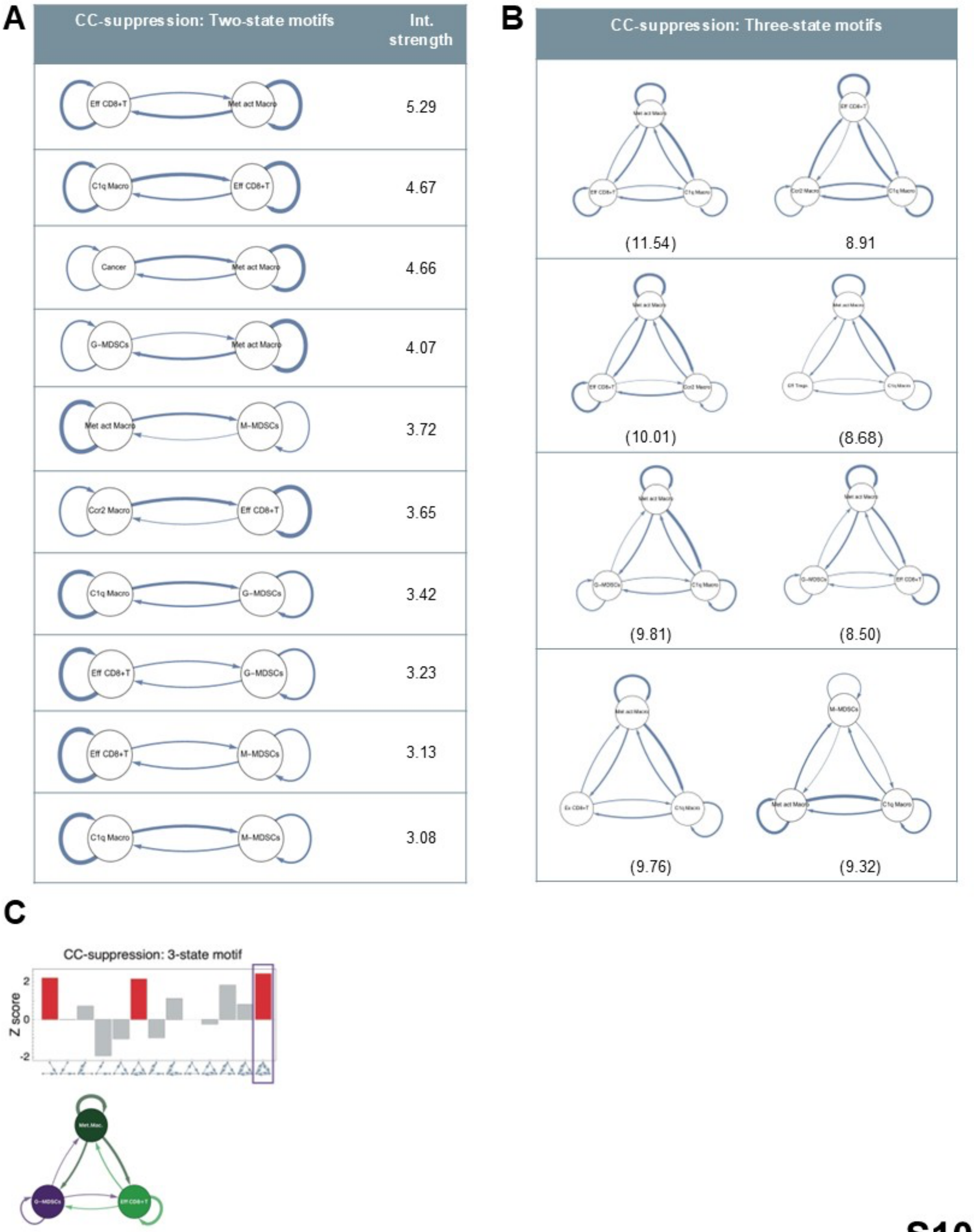
A
CC-suppression: Two-state motifs
Int. strength
Eff CD8+T
Met act Macro
5.29
C1q Macro
Eff CD8+T
4.67
Cancer
Met act Macro
4.66
G-MDSCs
Met act Macro
4.07
Met act Macro
M-MDSCs
3.72
Ccr2 Macro
Eff CD8+T
3.65
C1q Macro
G-MDSCs
3.42
Eff CD8+T
G-MDSCs
3.23
Eff CD8+T
M-MDSCs
3.13
C1q Macro
M-MDSCs
3.08
B
CC-suppression: Three-state motifs
(11.54)
8.91
(10.01)
(8.68)
(9.81)
(8.50)
(9.76)
(9.32)
C
CC-suppression: 3-state motif
Z score
2
0
-2

S10

A

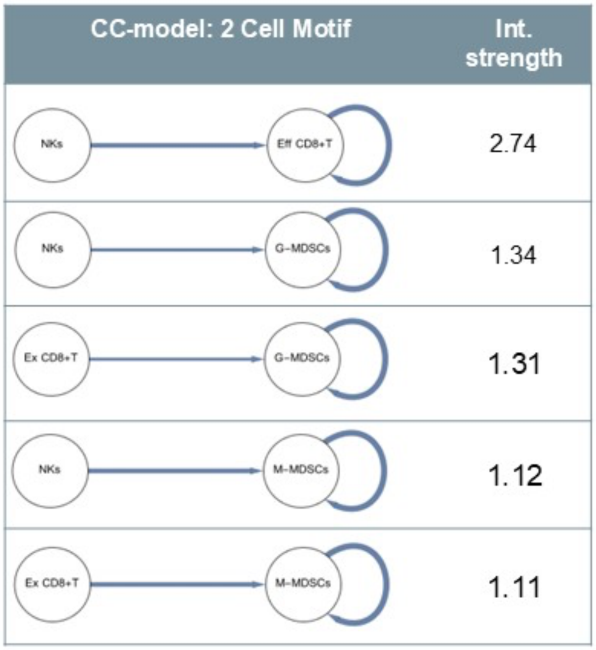


B

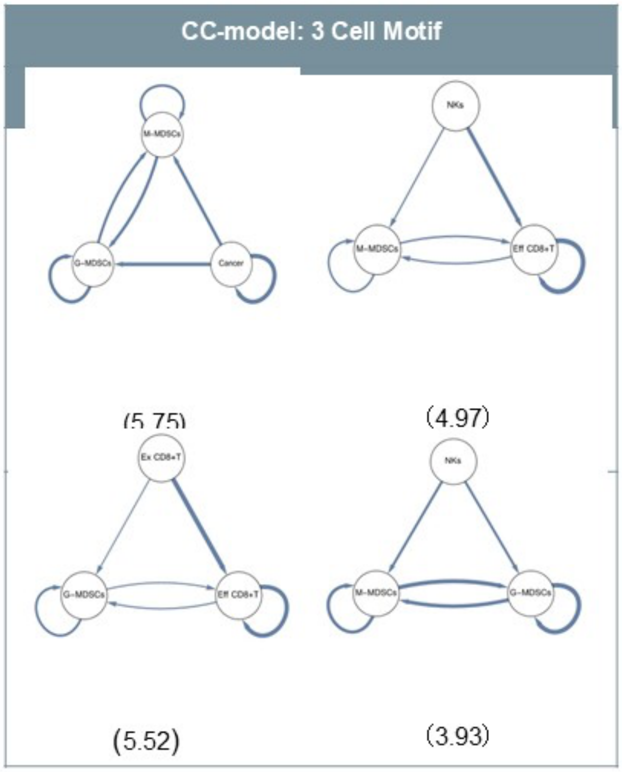


C

| Pathway | Log2fc E | Log2fc EPC | Log2fc PC |
|---|---|---|---|
| CCL | -0.64 | -0.43 | 1.12 |
| CD80 | -1.72 | off with EPC | -1.43 |
| CD86 | 0.30 | -0.94 | 0.78 |
| FN1 | -0.70 | 0.71 | 0.62 |
| GALECTIN | 0.03 | -1.52 | -0.01 |
| ICAM | 0.10 | -0.74 | -0.16 |
| ITGAL-ITGB2 | -1.49 | -1.53 | 0.11 |
| JAM | 0.83 | 0.60 | 0.69 |
| MHC-1 | -1.75 | -0.78 | 0.18 |
| SPP1 | -0.92 | 0.33 | -0.06 |
| THBS | off with E | off with EPC | off with PC |

D

| Pathway | Log2fc E | Log2fc EPC | Log2fc PC |
|---|---|---|---|
| CCL | -0.50 | -0.39 | 1.28 |
| GALECTIN | -0.37 | Off with EPC | 0.39 |
| ITGAL-ITGB2 | -1.12 | -1.30 | 0.48 |
| MHC-I | -1.86 | -0.98 | 0.20 |

E

| Pathway | Log2fc E | Log2fc EPC | Log2fc PC |
|---|---|---|---|
| ANNEXIN | -0.02 | -0.36 | 0.15 |
| BST2 | 0.48 | -0.04 | 0.71 |
| CCL | 0.03 | -0.26 | 0.20 |
| COMPLEMENT | 0.50 | -0.27 | -1.03 |
| CXCL | 4.45 | 4.40 | 5.14 |
| FN1 | -0.30 | 0.13 | 0.38 |
| GALECTIN | -0.03 | -1.62 | 0.15 |
| ICAM | 0.82 | -0.80 | -3.37 |
| JAM | 0.93 | 0.19 | 0.70 |
| LAIR1 | -1.82 | -1.91 | -0.88 |
| SELPLG | on with E | 0.00 | 0.00 |
| SPP1 | -0.57 | -0.30 | -0.56 |
| THBS | off with E | off with EPC | off with PC |

S11

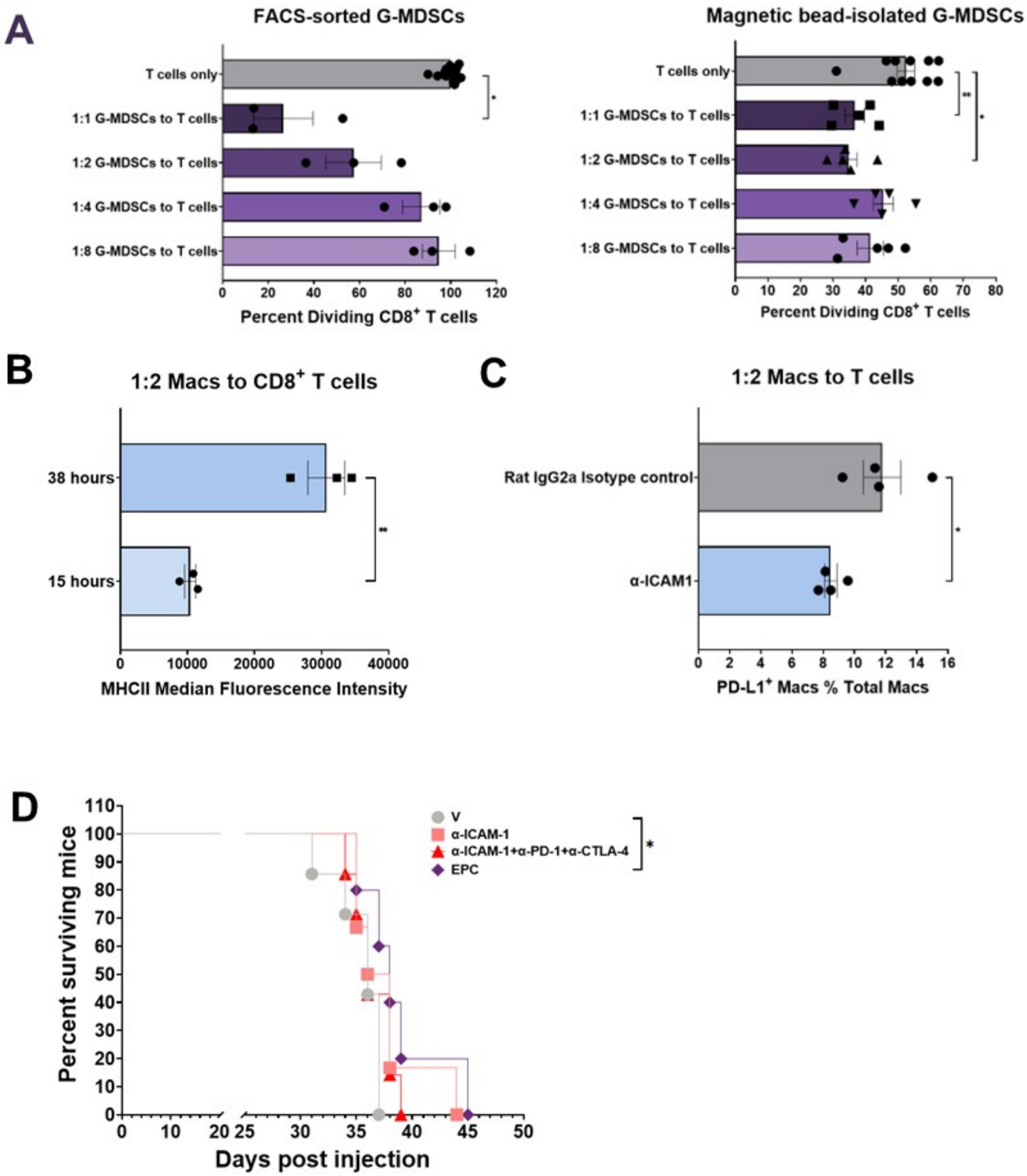
A
FACS-sorted G-MDSCs
T cells only
1:1 G-MDSCs to T cells
1:2 G-MDSCs to T cells
1:4 G-MDSCs to T cells
1:8 G-MDSCs to T cells
Percent Dividing CD8+ T cells
Magnetic bead-isolated G-MDSCs
B
1:2 Macs to CD8+ T cells
38 hours
15 hours
MHCII Median Fluorescence Intensity
C
1:2 Macs to T cells
Rat IgG2a Isotype control
α-ICAM1
PD-L1+ Macs % Total Macs
D
Percent surviving mice
Days post injection
V
α-ICAM-1
α-ICAM-1+α-PD-1+α-CTLA-4
EPC

S12

**A**

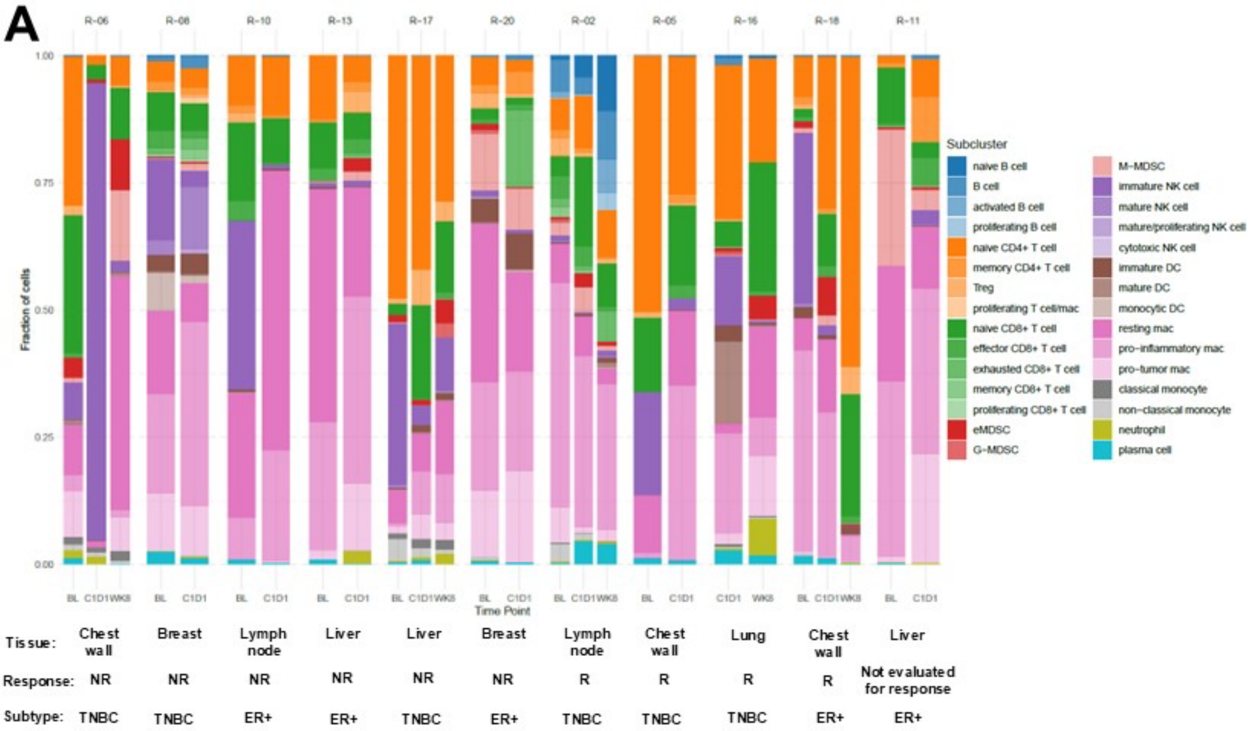


**B**

| Immune cell marker |
|---|
| CD45 |
| **Myeloid cell markers** |
| CD68 |
| CD33 |
| CD16 |
| **Lymphoid cell markers** |
| CD3 |
| CD20 |
| CD57 |
| **Cell subtype markers** |
| CD4 |
| CD8 |
| CD163 |
| CD15 |
| CD14 |
| DC-LAMP |
| DC-SIGN |
| Tox-Tox2 |
| FOXP3 |
| CD45RO |
| CD45RA |
| **Stromal markers** |
| Collagen |
| VIM |
| Podoplanin |
| αSMA |
| **Epithelial markers** |
| E-cadherin |
| CK (Pan cytokeratin) |
| **General markers** |
| Ruthenium 1 |
| Ruthenium 2 |
| Iridium 1 |
| Iridium 2 |
| Plasma membrane 4 |
| **Functional markers** |
| Ki-67 |
| PD-1 |
| PD-L1 |
| LAG3 |
| CD86 |
| pSTAT3 |
| Arginase 1 |
| Granzyme B |
| CD137 |
| HLA-DR |

**C**

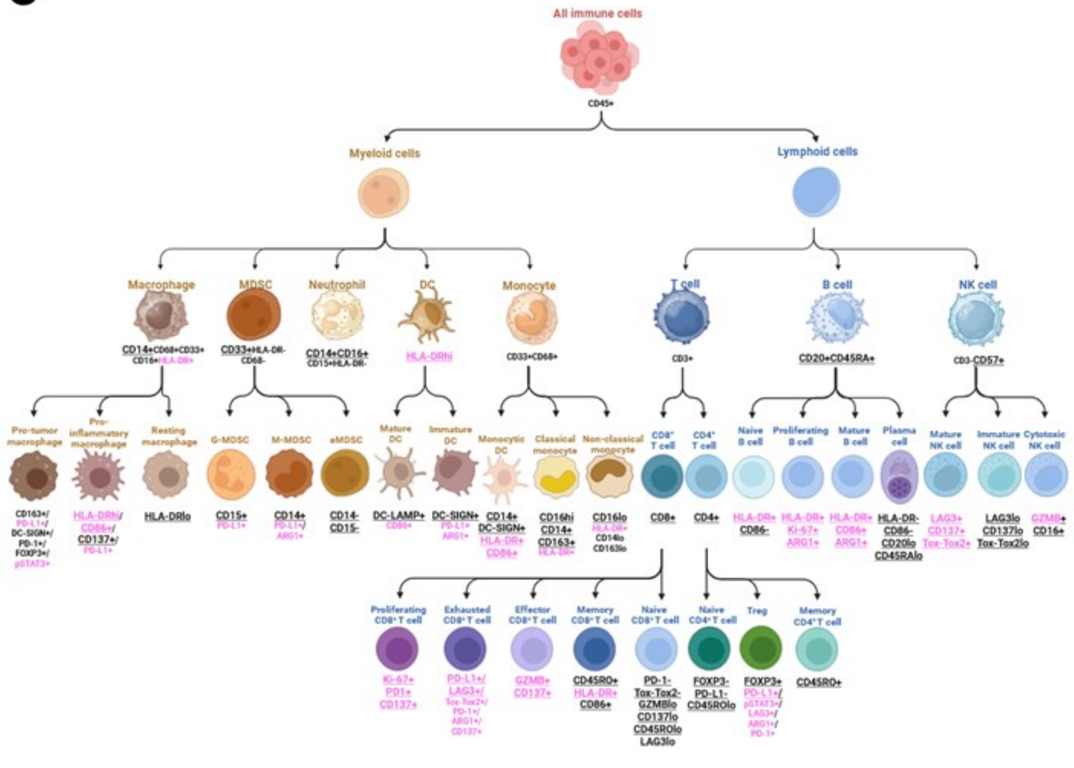


S13

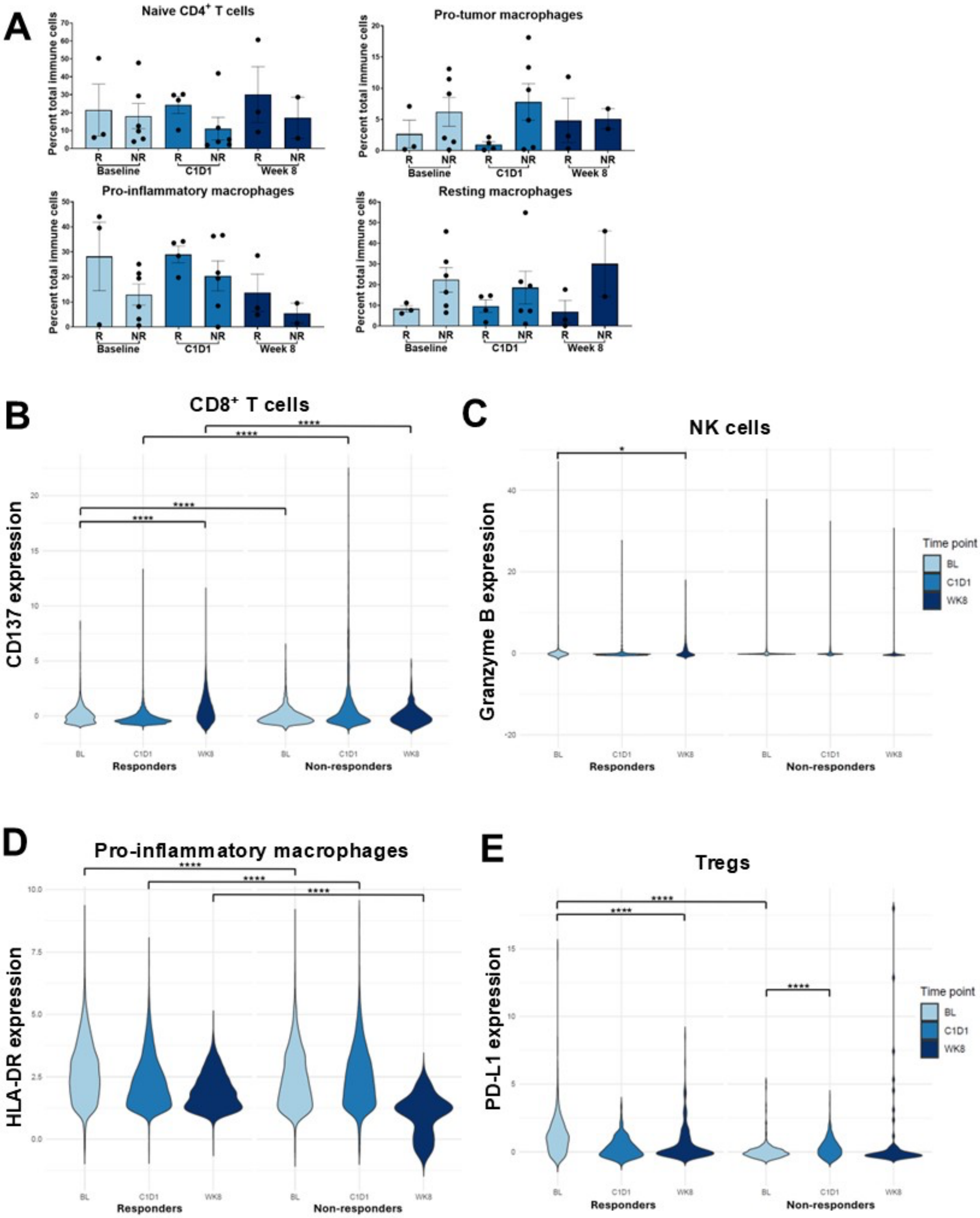


S14

INTERNAL

**Figure S1. Expression of fibroblast markers in mammary tumor CAFs and lipofibroblasts from lung metastases.** Dot/violin plot showing the expression of signature marker genes for cancer-associated fibroblasts (CAFs), lipofibroblasts (lipof.), and myofibroblasts (mCAFs) in CAFs identified in NT2.5 mammary tumors and lipofibroblasts identified in NT2.5-LM lung metastases.

**Figure S2. Flow cytometry and scRNA-seq analysis of mature myeloid cell proportions in NT2.5-LM lung metastases with treatments. A-C.** Dot plots showing the expression of signature marker genes for mature monocyte/macrophage **(A)**, neutrophil **(B)**, and dendritic cell **(C)** populations identified in NT2.5-LM lung metastases using the gene list in supplementary table 1 in the scRNA-seq dataset. **D.** Stacked bar plots with cell proportions per treatment for mature myeloid cell populations and a table with significant differences in cell proportions with treatments. **E.** Flow cytometry results for analysis of proportions of monocyte, dendritic cell, and macrophage populations identified via scRNA-seq in NT2.5-LM lung metastases with treatments (n=4-5/group), shown as a percentage of total single, live cells. Cell markers used for the gating of each population were designed based on gene expression observed via scRNA-seq in each cell population. Antibody panels used for flow cytometry are found in Supplementary File 2. V = vehicle, E = entinostat, P = anti-PD-1, C = anti-CTLA-4. Statistically significant differences between cell proportions in panel D were determined by analysis of false discovery rates (FDR), as determined by the propeller method included in the speckle R package162.

**Figure S3. Flow cytometry and scRNA-seq analysis of T cell proportions in NT2.5-LM lung metastases with treatments. A-C.** Dot plots showing the expression of signature marker genes for CD8+ T cell **(A)**, CD4+ T cell **(B)**, and other lymphoid populations and basophils **(C)** identified within the T cell major cluster observed in NT2.5-LM lung metastases in Figure 1C. Full gene list included in supplementary table 1. Statistically significant differences between cell proportions were determined via analysis of false discovery rates (FDR). **D.** UMAPs demonstrating the unsupervised projection of cell labels for T cell subsets identified using the projectTILs program on RStudio (top) in comparison with our manually annotated cell labels based on literature review of signature gene markers for T cells (bottom)**. E.** Stacked bar plots with cell proportions per treatment for T cell populations and a table with significant differences in cell proportions across treatments. **F.** Flow cytometry results for proportions of select T cell populations identified via scRNA-seq in NT2.5-LM lung metastases with treatments (n=4-5/group), shown as a percentage of total single, live cells. Cell markers used for the gating of each population were designed based on gene expression observed via scRNA-seq in each cell population. Antibody panels used for flow cytometry are found in Supplementary File 2. V = vehicle, E = entinostat, P = anti-PD-1, C = anti-CTLA-4. 7

**Figure S4. Determination of mature myeloid cell proportions with treatments in 4T1 lung metastases via flow cytometry.** Flow cytometry results for analysis of mature myeloid cell proportions, shown as a percentage of total single, live cells with treatments in 4T1 lung metastases (n=5-6/group). Cell markers used for the gating of each population were designed based on gene expression observed via scRNA-seq in each cell population. *p<0.05. Antibody panels used for flow cytometry are found in Supplementary File 2. V = vehicle, E = entinostat, P = anti-PD-1, C = anti-CTLA-4. Statistically significant differences were determined using the Kruskal-Wallis test.

**Figure S5. Determination of T cell proportions with treatments in 4T1 lung metastases via flow cytometry.** Flow cytometry results for analysis of T cell proportions, shown as a percentage of total single, live cells with treatments in 4T1 lung metastases (n=5-6/group). Cell markers used for the gating of each population were designed based on gene expression observed via scRNA-seq in each cell population. Antibody panels used for flow cytometry are found in Supplementary File 2. *p<0.05. V = vehicle, E = entinostat, P = anti-PD-1, C = anti-CTLA-4. Statistically significant differences were determined using the Kruskal-Wallis test.

**Figure S6. Analysis of MDSC and NK cell populations in NT2.5-LM lung metastases with treatments by scRNA-seq. A.** UMAP of subclusters identified in the myeloid-derived suppressor cell (MDSC) major cluster (Figure 1C) in NT2.5-LM lung metastases via scRNA-seq. **B.** Dot plot showing the expression of signature markers for monocytic MDSCs (M-MDSCs) and granulocytic MDSCs (G-MDSCs) identified in A. **C.** Stacked bar plots showing MDSC proportions with treatments and accompanying table with statistically significant differences identified between treatment groups, determined via an unpaired t-test. **D-E.** Flow cytometry analyses of NT2.5-LM **(D)** and 4T1 **(E)** lung metastases showing proportions of M-MDSCs and G-MDSCs as a percentage of total single, live cells with treatments. **F.** UMAP of subclusters identified in the NK cell major cluster (Figure 1C) in NT2.5-LM lung metastases via scRNA-seq. **G.** Dot plot showing expression of signature markers for the three NK cell populations identified in **(F)**. **H.** Stacked bar plots showing the proportion of NK cell populations with treatments via scRNA-seq. **I.** Flow cytometry analyses of NT2.5-LM lung metastases showing proportions of NK cell populations as a percentage of total single, live cells. Cell markers used for the gating of each population were designed based on gene expression observed via scRNA-seq. Antibody panels used for flow cytometry are found in Supplementary File 2. V = vehicle, E = entinostat, P = anti-PD-1, C = anti-CTLA-4. Statistically significant differences were determined using the one-way ANOVA for panels D and I. *p<0.05. n=4-5/group for D, I, and n=5-6/group for E.

**Figure S7. Cell chat analysis of B cells in NT2.5-LM lung metastases with treatments and analysis of B cell proportions and expression of functional markers in patient tumors via IMC. A.** UMAP showing the expression of *Itgam* (CD11b) in B cell subclusters (left) along with UMAP of B cell subclusters (right). **B.** Heatmap showing differential interaction strengths for all ligand-receptor pairs considered in the cell chat analysis between vehicle (V) and entinostat (E)+ anti-PD-1 (P) + anti-CTLA-4 (C) treatment. All cell populations shown were included in the cell chat analysis. Cells that participate as the source of signaling are shown on the left column, while cells that receive signals are shown on the bottom. Bars next to and above the heatmap represent the differential interaction strength for all ligand-receptor pairs that the cell type participates in. **C-D.** Bar graphs depicting interaction strength between *Cd274* (PD-L1) on Bregs and *Pdcd1* (PD-1) on exhausted CD8+ T cells **(C)** and between *Cd86* on mature B cells 2 and *Cd28* on primed Th2 cells **(D)** in NT2.5-LM lung metastases with treatments as determined by cell chat analysis. **E.** Flow cytometry analysis of NT2.5LM cells (gated as CD45-HER2+ single, live cells) treated with serum from either healthy mice or mice bearing NT2.5-LM lung metastases treated with V, E, or EPC (n=2-3/group). PE-conjugated anti-mouse IgG was used to detect levels of NT2.5-LM targeting IgG in serum via quantification of median fluorescence intensity. Antibody panels used for flow cytometry are found in Supplementary File 2. **F.** Proportions of plasma cells in patient tumors stratified by treatment response and time point (n=2-6/group) as determined by imaging mass cytometry (IMC) analysis. *p<0.05. R = responder. NR = Non-responder, C1D1 = post-entinostat, week 8 = post entinostat + anti-PD-1 + anti-CTLA-4. **G.** Proportions of plasma cells, activated B cells, and proliferating B cells in all IMC patient samples across all time points containing a TLS or germinal center (lymph node samples) in at least 1 region of interest (ROI; n=7) compared to samples without a TLS or germinal center (n=19). **p<0.01. **H.** Visualization of IMC masks on patient tumor images showing the assignment of each cell category represented as a different color for patients R-06, R-08, R-10, R-02, and R-16 (left to right) at baseline, C1D1, and week 8 timepoints. White rectangles within images outline tertiary lymphoid structures (TLS) or germinal centers (in lymph nodes). Samples were not obtained for patients R-08 and R-10 at week 8 and R-16 at baseline. A TLS was defined as an aggregation of 8 or more B cells/plasma cells within a 125 μm radius in non-lymph node tissues. **I.** Expression of LAG3 and HLA-DR in mature B cells and CD137 in total B cells from tumors in responders (n=4) compared to non-responders (n=6). ****p<0.0001. Statistically significant differences were determined using the Mann-Whitney test for panel H, the unpaired t-test for panel I, and one-way ANOVA for panel E.

**Figure S8. Cancer cell heterogeneity and treatment effect on cancer cells. A.** Umap of sub-states in cancer cells within breast-to-lung metastases. **B.** The proportion of cells in each sub-state by treatment condition. **C.** Top 10 marker genes for each cancer cell state. **D.** Umap of the cell cycle phase of cancer cells. **E.** Proportion of cells in each cell cycle phase by treatment condition. **F.** Linear discriminant analysis (LDA) embedding quantifies treatment effect size. **G.** Top 10 genes explaining treatment difference identified by Wilcoxon rank sum test (DE) and linear discriminant analysis (LDA).

**Figure S9. Treatment effect on cancer cells. A.** Expression of top 10 genes explaining treatment differences identified by logistic regression (sLR) and iterative logistic regression (iLR). **B.** UCell was applied to score stemness genes in cancer cells by treatments, and the difference in scores between vehicle and other treatments was tested by t-test. **C.** UCell was applied to score mesenchymal genes in cancer cells by treatments, and the difference in scores between vehicle and treatments was tested by t-test. **D.** UCell was applied to score PAM50 Myoepithelial (left) and Basal (right) genes on cancer cells by treatments, and the difference in scores between vehicle and treatments was tested by t-test. (*p<0.05; **p<0.01).

**Figure S10. CC-suppression cell circuit analysis. A.** CC-suppression most enriched two-node motif instances ranked by total strength. **B.** CC-suppression most enriched three-node motif instances, ranked by total strength. **C.** The most enriched three-cell motif and a particular instance of the top 3-state motifs selected from CC-suppression.

**Figure S11. CC-model cell circuit analysis. A.** CC-model most enriched two-node motif instances ranked by total strength. **B.** CC-model most enriched three-node motif instances ranked by total strength. **C.** Top changes in interaction strengths with treatment for pathways involving signaling from metabolically activated macrophages to effector CD8+ T cells, with the log fold change (Log2fc) due to treatment. **D.** Top changes in interaction strengths with treatment for pathways involving signaling from G-MDSCs to Effector CD8+ T cells. **E.** Top changes in interaction strengths with treatment for pathways involving signaling from Metabolically activated macrophages to G-MDSCs. Top pathways appearing in multiple comparisons were highlighted in blue.

**Figure S12. Flow cytometry analyses of CD8+ T cell co-cultures with G-MDSCs/macrophages from NT2.5-LM lung metastases. A.** Proportion of CD8+ T cells undergoing proliferation when either cultured alone or in co-culture with G-MDSCs obtained from NT2.5-LM lung metastases via FACS sorting of single, live CD45+CD11b+F4/80-Ly6G+Ly6C+ cells (left) or magnetic-bead pull-down (right) at ratios of 1:1, 1:2, 1:4, and 1:8 G-MDSCs to T cells (n=3-5/group). *p<0.05, **p<0.01, as determined via the Kruskal-Wallis test. **B.** Expression of MHC-II by median fluorescence intensity in total macrophages isolated from NT2.5-LM lung metastases and immediately evaluated (15 hours) versus stored overnight at 4 °C in Miltenyi MACS® Tissue Storage Solution (38 hours). n=3/group. **C.** Expression of PD-L1 in total macrophages co-cultured with CD8+ T cells treated with anti-ICAM-1 or Rat IgG2a isotype control (n=4) as determined via median fluorescence intensity quantification of PD-L1 using flow cytometry. Statistically significant differences were determined using the unpaired t-test for panels B-C. Co-culture assays illustrated in panels A-C were conducted as described in Figure 5. Antibody panels used for flow cytometry are found in Supplementary File 2. **D.** Survival experiment using the NT2.5-LM experimental model of lung metastasis (n=5-7/group). NeuN mice were injected into the tail vein with 100,000 NT2.5-LM cells and treated with vehicle (V) or 5 mg/kg entinostat (E) 5 times/week, 100 µg/dose anti-PD-1 twice/week (P), 100 µg/dose anti-CTLA-4 twice/week (C), or 100 µg/dose anti-ICAM-1 5 times/week starting on day 17 post-injection of NT2.5-LM cells. Statistically significant differences were determined using the log-rank test for survival.

**Figure S13. Proportions of immune cells with treatments in all patients and the IMC protocol used for analysis of immune cells in patient tumor samples. A.** Stacked bar plots showing the proportions of all immune cell populations identified via imaging mass cytometry (IMC) analysis of tumor samples from patients by treatment timepoint, with sample characteristics described underneath. BL = baseline, C1D1 = post-entinostat, WK8 = post-entinostat + anti-PD-1 + anti-CTLA-4 (week 8 timepoint), R = responder, NR = non-responder, TNBC = triple-negative breast cancer, ER+ = estrogen receptor-positive. **B.** Table of IMC general markers and cell-specific markers included in the analysis categorized by immune cell markers, myeloid cell markers, lymphoid cell markers, cell subtype specific markers, stromal cell markers, epithelial cell markers, functional markers, and general markers. **C.** Hierarchy of immune cell populations showing signature cell markers for populations of cells identified in high-level clusters (3$^{rd}$ level from the top) and immune cell subpopulations (4$^{th}$-5$^{th}$ levels from the top). Cell markers demonstrating characteristic expression in that cell type that were identified in every subcluster within that cell population are denoted by underlined text, and functional markers are denoted by magenta-colored text.

**Figure S14. Differences in immune cell proportions and expression of functional markers associated with treatments in patient IMC data. A.** Proportions of naïve CD4+ T cells, pro-inflammatory macrophages, pro-tumor macrophages, and resting macrophages in patient tumors stratified by treatment response and timepoint (n=2-6/group), as determined using imaging mass cytometry (IMC). **B-E.** Expression of CD137 in CD8+ T cells (**B**), granzyme B in total NK cells (**C**), HLA-DR in pro-inflammatory macrophages (**D**), and PD-L1 in Tregs (**E**) from tumors in responders (n=4) compared to non-responders (n=6). *p<0.05, ****p<0.0001, as determined using unpaired T tests for panels B-E.
